# Probiotic formula intervention in infants leads to colonization and competitive strain displacement independent of IgA binding

**DOI:** 10.64898/2026.06.03.729775

**Authors:** Michael Vig Merino, Svenja Weißenberger, Anastasia Ermolova, Elise Dhilly, Roan Spadazzi, Lior Levy, María Carmen Collado, Tomer Hertz, Hélène Omer, Vitor Heidrich, Nicola Segata, Martin Larsen, Melanie Schirmer, Dirk Haller

**Affiliations:** Chair of Nutrition and Immunology, Technical University of Munich, Gregor-Mendel-Str. 2, 85354 Freising, Germany; Professorship for Translational Microbiome Data Integration, Technical University of Munich, Gregor-Mendel-Str. 2, 85354 Freising, Germany; Center for Organoid Systems, Technical University of Munich, Garching, Germany; Sorbonne University, Inserm UMR-S1135, Centre d’Immunologie et des Maladies Infectieuses (CIMI Paris), 91 boulevard de l’Hopital, 75013 Paris, France; Dept. CIBIO, University of Trento, 38123, Italy; Dept. of Microbiology, Immunology and Genetics, Ben-Gurion University of the Negev, Beer-Sheva, 84105, Israel; Dept. of Biotechnology, Institute of Agrochemistry and Food Technology, Spanish National Research Council (IATA-CSIC), 46980 Paterna, Valencia, Spain; ZIEL Institute for Food and Health, Technical University of Munich, Germany

**Keywords:** Microbiome, infant, shotgun metagenomics, Bifidobacteria, GOS, formula, breast-feeding, strain diversity, microbial transmission, IgA response

## Abstract

Early-life microbiota assembly is affected by environmental exposure, diet, and immunity. We performed shotgun metagenomics sequencing of fecal samples (N=401) from a controlled infant intervention trial to evaluate how pre- and probiotic exposure affects bacterial strain dynamics, maternal-to-offspring strain transmissibility, strain persistence, and host immune responses. Species- and strain-level analyses revealed that the early-life intervention with *Bifidobacterium longum* subsp. *infantis (BL* subsp. *infantis), Bifidobacterium breve,* and galacto-oligosaccharides (GOS) replaced main resident strains of these species unlike breast-feeding and exclusive infant formula. *BL* subsp. *infantis* low-diversity clusters were representative of several populations and were enriched for genes linked to recombination, mobilization, and antimicrobial activity. lnterestingly, *Bifidobacterium bifidum* but not *Bifidobacterium longum* or *B. breve* was heavily targeted by mucosal IgA, regardless of feeding type. These findings suggest that early-life pro- and prebiotic interventions promote colonization of specific dominant strains, potentially by outcompeting and displacing existing strains, independently of host immune responses.

## lntroduction

Colonization of the infant gut microbiota starts at birth, and is largely driven by delivery mode^1–6^, dietary exposure^1,5,7^, priority effects^8–10^, vertical transmission^11–14^, and horizontal transmission^15^. It is well known that the infant microbiome shifts towards parental composition over time^16–18^. Interestingly, different feces metabolome profiles have been observed between formula-fed infants and breast-fed infants^19,20^ and in feces taxonomic composition^18^ in early-life.

Key representatives of infant gut commensals include the genera *Bifidobacterium*, *Bacteroides*, *Veillonella,* and *Escherichia*. Bifidobacteria are dominant in early-life^9,19^ and are frequently used as probiotic supplementation^21^. Some species and subspecies have been linked to an early dominance and high prevalence (e.g., *Bifidobacterium breve*)^8^. In contrast, *Bifidobacterium longum* subsp. *infantis (BL* subsp. *infantis)* has been shown to be depleted in the first 2 months of life in high-income countries (HICs)^22^, but not in low- and middle-income countries (LMICs)^23^, hence, further studies are needed to characterize species colonization dynamics. Furthermore, as individual-specific members of the human microbiome, strains exhibit considerable genomic and functional heterogeneity within species^23,24^. Accurate strain-level analyses are therefore critical to elucidate the potential interplay between dietary interventions, age, and strain tracking at the taxonomic and functional level.

Immunoglobulin A (IgA) is the main antibody in the gut^25^, with most antibody-secreting plasma cells located in the intestine to sustain continuous mucosal secretion of secretory IgA^26^. IgA helps maintain gut microbial balance by modulating microbial colonization dynamics and host immune responses^27,28^. The development of the infant microbial ecosystem is essential for shaping the host immune system. Early-life crosstalk between the gut microbiota and the immune system influences long-term host health, and it is partly regulated by IgA derived from maternal human milk and increasingly from the infant itself^27^. Impaired IgA responses are linked to higher infection risk, autoimmune disease, and bacterial translocation across the intestinal barrier^29–32^.

We recently performed a controlled intervention trial with an infant cohort (n = 210 participants) during their first year of life (Infantibio-II: German Clinical Trial DRKS00012313) in which we compared a breast-fed reference group against four randomized formula-fed groups: a placebo (formula A), combined with probiotics (supplemented with bifidobacteria; formula B), prebiotics (supplemented with galacto-oligosaccharides (GOS); formula C) and synbiotic (supplemented with both bifidobacteria and GOS; formula D)^19^. By performing 16S rRNA-based microbiota profiling of longitudinally sampled infant feces (n = 998), we found that age, but not dietary exposure, was the primary driver of microbiota composition. Untargeted metabolomics analysis revealed that the exposure to either breast milk or infant formula was associated with different metabolite profiles.

In this study, we analyzed how species- and strain-level dynamics, particularly those of early-and late-dominant colonizers, together with IgA responses, parental-to-offspring transmissibility, and strain persistence, are influenced by age and exposure to pre- and probiotics. Shotgun metagenomics was performed using a standardized pipeline which combined two different approaches: a reference-based approach and a de-novo assembly-based approach.

## Results

### *Bifidobacterium* dominates the first months of life, but species dynamics is significantly impacted by early probiotics exposure

We analyzed a longitudinal dataset of 401 fecal shotgun metagenomes (obtained from N = 60 infants, n=297 samples; and their paired parents, n= 104) which included five infant time points (1, 3, 7, 12, and 24 months of age). We first assessed patterns of bacterial alpha diversity (richness and Shannon index) and beta diversity (Bray-Curtis dissimilarity). Diversity patterns were consistent with those previously observed using 16S rRNA gene amplicon sequencing^19^. As expected, infant samples of the latest sampling time point (24-months of age) distinctively clustered towards the parents (data not shown) and away from the first-year samples, showing higher values for richness and Shannon diversity index (**Figure 1A**, **Figure S1A**).

**Figure 1.**
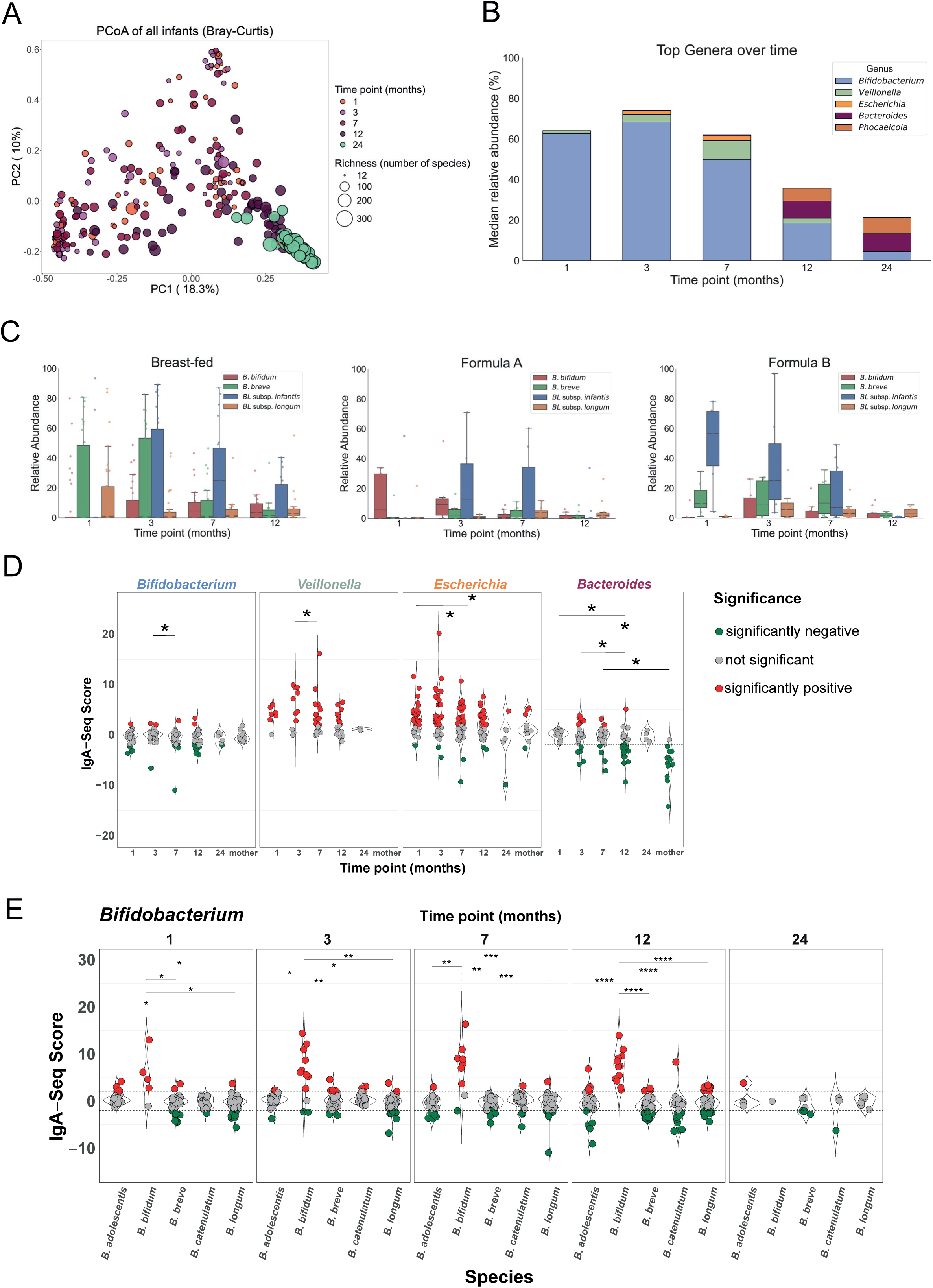
Effect of age and feeding mode on the infant microbial gut composition, lgA-binding levels and *Bifidobacterium* species growth dynamics. (A) PCoA plot of beta-diversity (Bray-Curtis) of all infant samples (n= 297) grouped according to infant age (months) with richness (alpha-diversity) shown by size. (B) Distribution over time of the five dominant genera in all infant fecal samples (n= 297), represented by median relative abundance (%). Dominant genera included those with the highest overall relative abundance across all infant samples: *Bifidobacterium*, *Veillonella*, *Escherichia*, *Bacteroides*, and *Phocaeicola*. (C) Top *Bifidobacterium* species by median relative abundance (%) for breast-fed infants (left), formula A-fed infants (placebo, middle) and formula B-fed infants (probiotics, right) with each color representing a species. (D) Median IgA-binding scores of the top genera over time over all infants and paired mothers (n = 135 samples). Significance is measured between the time points for each genus (paired Wilcoxon test). Stars indicate p-values significance (with Benjamini-Hochberg correction): p<0.05 (*). (E) Median IgA-binding scores of *Bifidobacterium* distributed by species over time. Significance is measured between the species for each time point (unpaired Wilcoxon test). Stars indicate p-values significance (with Benjamini-Hochberg correction): p<0.05 (*), p<0.01 (**), p<0.001 (***), p<0.0001 (****), p<0.00001 (*****)

To investigate whether feeding or age influenced the species-level microbiome composition in infants, we performed PERMANOVA on Bray-Curtis dissimilarities. The model included the interaction of feeding and time point (age) while adjusting for potential confounders (sex, delivery mode, and maternal antibiotic exposure). To account for the non-independence of repeated samples from the same infants, permutations were stratified by infant ID using a blocked permutation design, and marginal tests were used for each term. A significant interaction between feeding and age was observed (p = 0.0015), indicating that the association between feeding and microbiome composition was age-dependent rather than uniform across time.

Exploratory pairwise PERMANOVA comparisons among feeding groups for all time points together followed by Benjamini-Hochberg (BH) correction^33^ revealed pronounced differences in overall Bray-Curtis dissimilarity involving feeding in the 24-months group (adj. p-value = 2.2e-04), consistently showing the largest effect sizes (R2 = 0.14–0.22). In addition to feeding, an analogous pairwise PERMANOVA analysis was performed for all age groups, revealing that all pairwise comparisons between time points were significant after BH correction (adj. p-value range = 1.4e-04-2.9e-04), indicating pronounced temporal differences in species level composition.

Because the interaction of feeding and age was significant, we further explored the effects of feeding within each time point using pairwise PERMANOVA. At month 1 of age, breast-fed samples were significantly different from formula groups B (probiotics, adj. p-value = 0.001) and D (synbiotic, adj. p-value = 0.001), while formula group A (placebo) significantly differed from groups B (adj. p-value = 0.026) and D (adj. p-value = 0.023). Pairwise comparisons between the feeding groups at 3, 7, and 12 months were not significant (adj. p-value > 0.05).

Multivariate dispersions differed significantly across feeding groups (betadisper, p = 1e-04) and across age groups (betadisper, p = 1e-04), indicating substantial heterogeneity in within-group community variability and warranting cautious interpretation of the pairwise PERMANOVA results.

The *Bifidobacterium* genus dominates in the first year of life, especially in the first month samples, and then quickly diminishes. In contrast, the relative abundances of late colonizers, such as *Bacteroides* and *Phocaeicola*, increased over time (**Figure 1B**). The median relative abundance of *Bifidobacterium* was the highest at 3 months of age (68.4%). At this time point, formula group D (synbiotic) revealed the highest median relative abundance (78.82%), followed by breast-fed (71.85%), and formula groups B (probiotics; 65.3%), C (prebiotics; 58.34%) and A (placebo; 52.78%) (**Figure S1B**). The lowest *Bifidobacterium* levels were observed at 24 months of age (4.42%).

*BL* subsp. *infantis* was strongly enriched at 1-month of age when infants were exposed to *BL* subsp. *infantis* supplemented formula B (probiotics) compared to control formula A (adj. p-value = 0.041) and to breast-feeding (adj. p-value = 7e-04) (**Figure 1C**). Early GOS exposure at month 1 (prebiotics/formula C; **Figure S1C**) was associated with a similar colonization pattern but did not reach significance (adj. p-value = 0.32 against placebo formula A and adj. p-value = 0.078 against breast-feeding). However, the effect was the strongest with the synbiotic combination (formula D; **Figure S1D**) (adj. p-value = 0.029 against formula A and adj. p-value = 4e-04 against breast-feeding). In sharp contrast, *B. breve* intake at 1 month of age did not support a higher colonization by *B. breve* (Kruskal Wallis^34^, p = 0.27) compared to any other feeding exposure, suggesting species-specific colonization patterns.

While these early, species-specific colonization effects were evident at 1 month of age, they did not appear to shape the broader microbiome at later time points. Taxonomic composition at the family level in infants at 24 months was highly similar to that of the parents (**Figure S1E**). Bray-Curtis dissimilarity showed minor differences in overall microbiome composition between feeding groups at 24 months of age (**Figure S1F**).

### Secretory lgA responses to *Bacteroides, Escherichia, Veillonella*, and *Bifidobacterium bifidum* are age-dependent and unaffected by diet or microbial abundance

We quantified host mucosal immune targeting of the microbiota by measuring endogenous secretory IgA (sIgA) coating of fecal bacteria using IgA-Seq technology. Briefly, bacteria were sorted into IgA-coated and non-coated fractions, and enrichment of each taxon in the IgA-positive fraction was expressed as a standardized IgA-Seq Z-score. Positive scores indicated enrichment in the IgA-coated fraction, whereas negative scores reflected enrichment in the non-coated fraction.

Using this metric, we observed that the bacterial genera *Veillonella* and *Escherichia* were heavily targeted by sIgA during the first year of life, with significantly elevated binding scores peaking at 7 and 3 months of age, respectively (**Figure 1D**). In contrast, IgA-Seq scores for *Bacteroides* progressively declined across the first year of life, approaching levels observed in the mothers by month 12. No significant association with age in sIgA reactivity was observed within the *Bifidobacterium* genus. However, among *Bifidobacterium* species, only *B. bifidum* displayed significantly high IgA-Seq scores across all infant time points in which they were detected (except month 24 where only one sample was profiled) (**Figure 1E**) independently of relative abundances (**Figure S2A**). Throughout the first year of life, *Veillonella*, *Escherichia* and *B. bifidum* were consistently and strongly targeted by IgA regardless of dietary exposure (**Figures S2B** and **S2C**). Interestingly, *B. longum,* which was one of the most abundant species during the first year of life, was not significantly targeted by IgA (**Figure 1E**).

### Probiotics exposure impacts the early life assembly of *Bifidobacterium* strain diversity

As observed in the phylogeny of *BL* subsp. *infantis* and *B. breve*, the supplementation with probiotics and prebiotics was associated with the clonal colonization and expansion of the selected dominant strain, potentially preventing naturally acquired strains related to breast-feeding and placebo formula-feeding from expanding (**Figure 2A** top left and top right). Two very distinct clusters were observed for these two species: a first high-diversity cluster defined as “cluster 1” (70 samples in *BL* subsp. *infantis* and 161 samples in *B. breve*), and a second low-diversity “cluster 2” which comprises the formula-contained strain (63 samples in *BL* subsp. *infantis* and 46 samples in *B. breve*). Conversely, *B. bifidum* (**Figure 2A** bottom left), *Veillonella parvula* (Figure 2A bottom right), *Bifidobacterium longum* subsp. *longum (BL* subsp. *longum)* (**Figure S3A**), *Escherichia coli* (**Figure S3B**), *Bacteroides uniformis* (**Figure S3C**) and *Phocaeicola vulgatus* (**Figure S3D**) did not show any impact related to dietary feeding exposure at the dominant strain level. No clear evidence of age-related effects was observed in the phylogenies of the analyzed species (**Figures S4A-F**).

**Figure 2.**
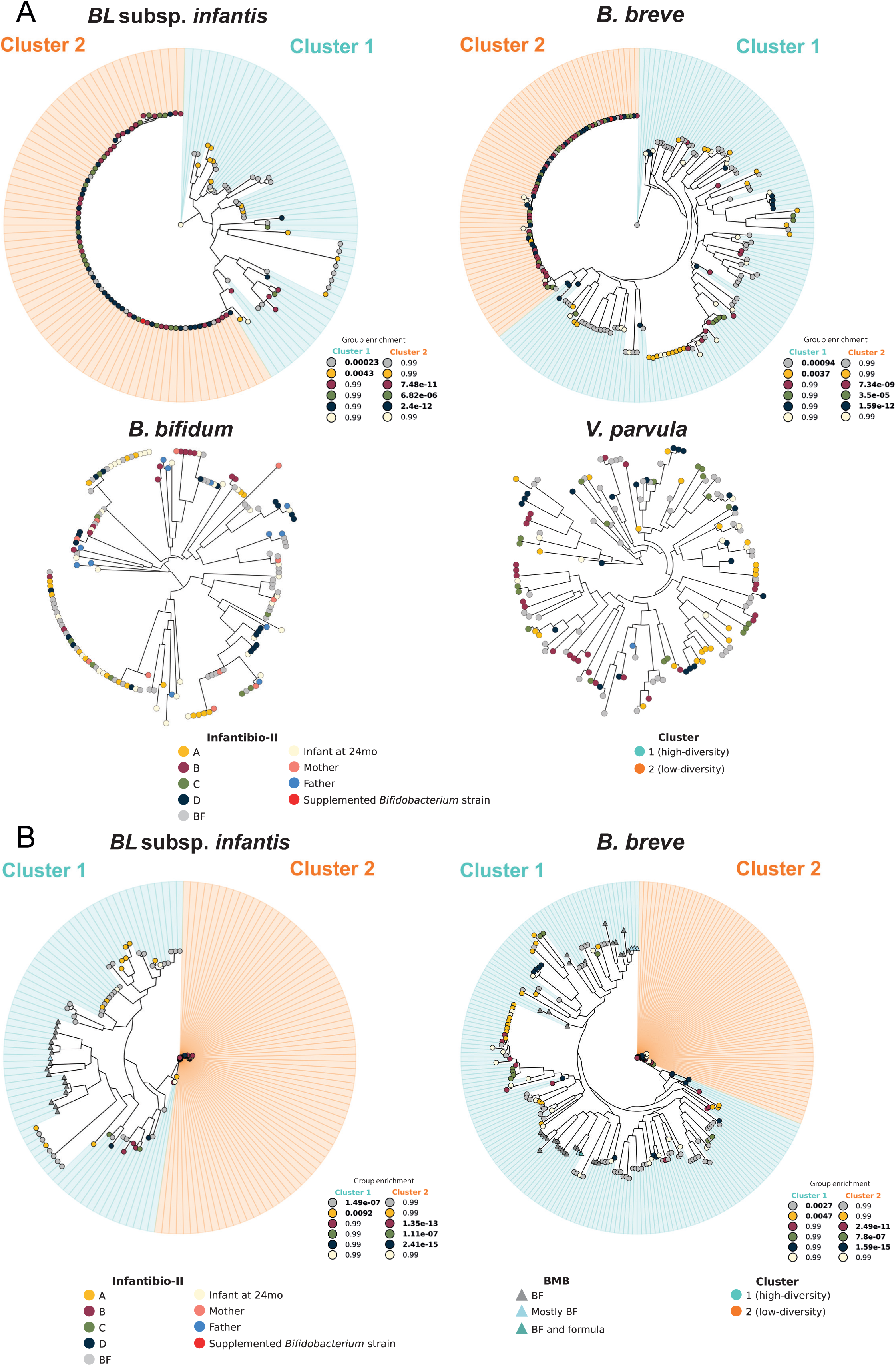
Effect of feeding mode in the *Bifidobacterium* dominant strain diversity. (A) Phylogenies of early infant colonizers, showing one dominant strain per sample (StrainPhlAn). The phylogeny includes 133 samples for *BL* subsp. *infantis*, 207 samples for *B. breve*, 168 samples for *B. bifidum*, and 148 samples for *V. parvula*, generated using 119, 200, 200, and 200 clade marker genes, respectively. Cluster 2 (low-diversity strains) was defined using strain identity thresholds relative to the reference probiotic strains, calculated from Kimura-corrected evolutionary distances (>96.02% for *BL* subsp. *infantis* and >97.28% *B. breve*). Feeding mode is indicated by color, with light red highlighting the reference supplemented *Bifidobacterium* strain. Adjusted p-values (with Benjamini-Hochberg correction) indicate significant over-representation of each feeding group (hypergeometric test, adj. p-value < 0.05) within clusters, according to the original group sizes. (B) Combined StrainPhlAn phylogenies of *BL* subsp. *infantis* and *B. breve* from the Infantibio-II cohort and BMB cohort, showing one dominant strain per sample. The phylogeny includes 152 samples for *BL* subsp. *infantis* and 243 samples for *B. breve*, generated using the same marker genes and parameters as described above. Reference supplemented *Bifidobacterium* strains are highlighted in light red. Identity thresholds were determined as specified above and were >96.84% for *BL* subsp. *infantis* and >97.55% for *B. breve*. Adjusted p-values (with Benjamini-Hochberg correction) indicate significant over-representation of each feeding group (hypergeometric test, adj. p-value < 0.05) within clusters, according to the original group sizes.

In the high-diversity cluster (cluster 1) of the *BL* subsp. *infantis* phylogeny, breast-fed and placebo (formula A) groups were overrepresented (adj. p-value = 2.3e-04 and 0.0043 respectively), while all supplemented formula groups (B, C and D) were enriched in the low-diversity cluster (cluster 2) (adj. p-value = 7.5e-11, 6.8e-06 and 2.4e-12 respectively) (**Figure 2A** top left). A similar pattern was observed with *B. breve*: breast-fed and formula A groups were significantly enriched in cluster 1 (adj. p-value *=* 9.4e-04 and 0.0037), while formula B, C and D groups were overrepresented in cluster 2 (adj. p-value *=* 7.3e-09, 3.5e-05, and 1.6e-12) (**Figure 2A** top right).

We additionally integrated longitudinal metagenomes from the Breast Milk Baby (BMB) cohort^22^ into our *BL* subsp. *infantis* and *B. breve* phylogenies. This cohort comprises paired stool and breast milk samples collected from 21 healthy mother-infant dyads (exclusively breast-fed, partially breast-fed, and formula-fed) during the first year of life. All of the recovered genomes (*BL subsp*. *infantis* n = 18; *B. breve* n = 35) fell into the high-diversity clusters (**Figure 2B**). While diversity was expected as most genomes were recovered from exclusively breast-fed infants (*BL* subsp. *infantis* n = 16; *B. breve* n = 30); all samples were grouped, suggesting a diverse bifidobacterial strain colonization in the absence of probiotic supplementation of the infant formula.

### *B. longum* and *E. coli* dominant strains in breast-fed infants are more similar to those found in their paired mothers compared to those in formula-fed infants

Three species were analysed for mothers-to-offspring transmissibility, including *BL* subsp. *longum* (**Figure S5A**), *E. coli* (**Figure S5B**), and *B. uniformis* (**Figure S5C**). *BL* subsp. *infantis*, *B. breve* and *V. parvula* were not detected in the overwhelming majority of parents, and the overall number of samples was too low for *P. vulgatus*. Sample sizes varied considerably between related and unrelated parent–infant pairs and across infant age for each species. Evolutionary distances from infants to related mothers were significantly lower than those from infants to unrelated mothers for *BL* subsp. *longum* (Kruskal Wallis, p = 2.8e-20), *E. coli* (Kruskal Wallis, p = 5.4e-04), and *B. uniformis* (Kruskal Wallis, p = 1.4e-23). Additionally, distances from infants to related fathers were also lower than those from infants to unrelated fathers for *BL* subsp. *longum* (Kruskal Wallis, p = 0.0080) and *B. uniformis* (Kruskal Wallis, p = 0.0015), but not for *E. coli* (Kruskal Wallis, p = 0.57). No significant difference was observed between infant sampling time points when analysing the distances from infants to related mothers for any of the three species. Distances to paired mothers were significantly impacted by dietary intervention for *BL* subsp. *longum* (Mann-Whitney, adj. p-value ≤ 0.01) and *E. coli* (Mann-Whitney, adj. p-value ≤ 0.01), with lower distances observed in breast-fed compared to formula-fed infants, but not for *B. uniformis* (Mann-Whitney, adj. p-value = 0.90). No significant difference was observed in the distances to related fathers in any of the analysed species: *BL* subsp. *longum* (Mann-Whitney, adj. p-value = 0.20), *E. coli* (Mann-Whitney, adj. p-value = 0.42), and *B. uniformis* (Mann-Whitney, adj. p-value = 0.69).

### *Bifidobacterium* shunt, glycogen degradation pathways and associated-genes related to carbohydrate breakdown are enriched in formula-fed infants

To elucidate gene pathways that differ by feeding type during the first year of life, we compared breast-fed and formula-fed samples (multivariate analysis with MaAsLin2). Pathways with adj. p-value < 0.25 were considered significant. The purine degradation pathway, a biomarker associated with disease risk^35^, was significantly enriched in formula-fed samples compared to breast-fed samples (adj. p-value = 5.5e-06). This signal was largely driven by *Enterococcus faecalis* (not shown). Other additional pathways were also enriched in the formula-fed group compared to breast-fed: L-glutamate and L-glutamine biosynthesis (adj. p-value = 0.0016), L-histidine degradation (adj. p-value = 0.0043), urea cycle (adj. p-value = 0.0145), beta-1,4 Mannan degradation (adj. p-value = 0.018), *Bifidobacterium* shunt (adj. p-value = 0.010), and glycogen degradation (adj. p-value = 0.056) (**Figure 3A**). Among these pathways, only the last two were strongly associated with bifidobacteria.

**Figure 3.**
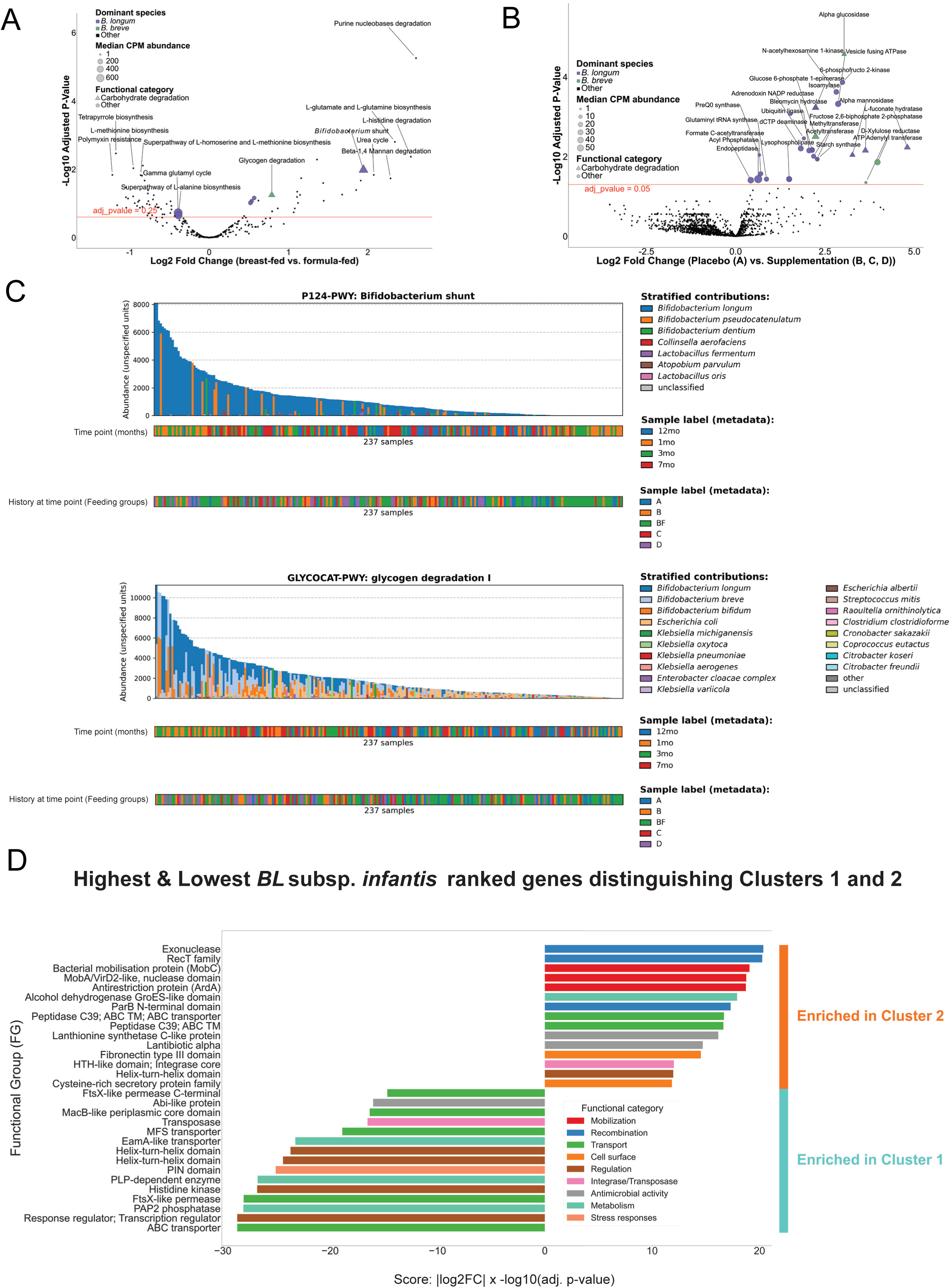
Functional profiles (gene families, pathways and PFAMs) according to feeding groups and clusters. (A) Volcano plot of 298 HUMAnN3 gene pathways (formula-fed against breast-fed as control). Pathways were highlighted by their Log2FC (coef) from MaAsLin2 linear model output. 73 pathways were significant; 225 were non-significant (threshold is adj. p-value < 0.25, highlighted as a red line). The median copies per million (CPM) abundance contribution (size) of each dominant *Bifidobacterium* species (color) were also shown. (B) Volcano plot of 2,688 detected gene families (HUMAnN3, supplemented formula groups B, C and D against placebo formula A as control). 42 gene families were significant; 2,646 were non-significant (MaAsLin2 linear model, adj. p-value < 0.05, threshold highlighted as a red line): Families were highlighted by their Log2FC (coef) from MaAsLin2 results, and each species contribution was shown as in Fig. 3A. (C) HUMAnN3 barplots of 237 first-year infant samples showing the abundances (in copies per million or CPM) of the *Bifidobacterium* shunt and glycogen degradation pathways stratified by the top bacterial contributors (indicated by color). (D) Top *BL* subsp. *infantis* genes with annotated Pfam domains significantly differed between cluster 1 and cluster 2, based on de-novo assembly–derived gene abundances and corresponding metagenomic species pangenome (MSP) (MaAsLin2 linear model, adj. p-value < 0.05). Genes with the highest scores are enriched in cluster 2, whereas those with the lowest scores are enriched in cluster 1.

The genes encoding 6-phosphofructo 2-kinase and glucose 6-phosphate 1-epimerase, related to the *Bifidobacterium* shunt pathway, a pathway unique to *Bifidobacterium*, were overabundant in the supplemented formula groups (B, C, and D), compared to the placebo (formula group A; adj. p-value = 1.4e-04 and adj. p-value = 0.0031, respectively) (**Figure 3B**). Several carbohydrate breakdown related genes were also enriched in the supplemented formula groups (B, C, and D) compared to the breast-fed group. These included alpha glucosidase (adj. p-value = 2.7e-05), which is involved in the degradation of human milk oligosaccharides (HMOs), and the debranching enzyme isoamylase (adj. p-value = 5.9e-04). Alpha mannosidase, another enzyme associated with HMOs breakdown, was also enriched in these formula groups compared to breast-fed infants (adj. p-value = 0.009).

While both bifidobacteria-related pathways are linked to the breakdown of carbohydrates, the *Bifidobacterium* shunt pathway was mostly dominated by *B. longum* (median CPM = 500.04) (**Figure 3C** top panel), while the glycogen degradation was primarily driven by *B. breve* (median CPM = 184.51) (**Figure 3C** bottom panel).

### *BL* subsp. *infantis* low-strain-diversity and high-strain-diversity clusters are characterized by different functional profiles

We then investigated the two *BL* subsp. *infantis* clusters (n = 133) in more detail using assembly-based analyses and were able to identify 7,942 genes that were unique to *BL* subsp. *infantis* Metagenomic Species Pangenome (MSP) 0009 (7,359 genes after 10% prevalence filtering). Of these genes, 2,605 were significantly different between the two clusters (adj. p-value < 0.05, MaAsLin2).

We next sought to elucidate which annotated de-novo assembled genes ranked as the most differentially abundant between the two clusters. For this purpose, we assigned a score as llog2FCl x -log10(adj. p-value) to rank these genes by significance and fold change (FC), so that those with positive scores were enriched in cluster 2 and those with negative scores were enriched in cluster 1. Only genes with predicted protein Pfam domains were considered. We observed that genes related to exonucleases (score = 20.35), the bacterial mobilisation protein (MobC) (score = 19.07), lanthionine synthetase (score = 16.16) and lantibiotics (score = 14.71) were significantly differentially prevalent in the *BL* subsp. *infantis* low-diversity cluster (cluster 2). In contrast, transport of solutes: ABC transporter (score = −28.65), Major Facilitator Superfamily (MFS) transporter (score = -18.86); response and transcription regulators (score = −28.65); lipid metabolism: PAP2 superfamily (score = −28.08); and stress responses: PIN-like domain (score = −25.07) were major functional groups found in the *BL* subsp. *infantis* high-diversity cluster (cluster 1) (**Figure 3D**).

### Low-diversity *BL* subsp. *infantis* strain clusters were also revealed within a global *Bifidobacterium* strain diversity framework

Comparing the strains from our infant samples against a collection of global *Bifidobacterium* reference genomes from 651 italian infant samples from the CM_MTB cohort^15^, 2,151 infant and children samples from cMD3 (from 14 countries, 9 westernized (Estonia, Finland, United Kingdom, India, Italy, Luxembourg, Russia, Sweden, United States) and 5 non-westernized (Ethiopia, Ghana, Peru, El Salvador, Tanzania); curatedMetagenomicData, version 3.18^36^), 76 food metagenomic samples from cFMD (curatedFoodMetagenomicData, version 1.2.1^37^). and 724 isolate reference genomes (MetaRefSGB, version vJan25^37,38^) enable comparative strain-level diversity analysis across four *Bifidobacterium* species: *BL* subsp. *infantis* (**Figure 4A**), *B. breve* (**Figure 4B**), *B. bifidum* (**Figure 4C**), and *BL* subsp. *longum* (**Figure 4D**). Low-diversity clusters associated with *BL* subsp. *infantis* and *B. breve* from our cohort were again revealed in this analysis. Importantly, low-diversity clusters were observed in infant cohorts of similar age in three of the four *Bifidobacterium* species (except *BL* subsp. *longum*). Addition of the isolate reference genomes to the phylogenies (**Figure 4**, green stars) showed that several commercial strains were located near low-diversity clusters in *BL* subsp. *infantis, B. breve*, and *B. bifidum*. Notably, several isolates fell close to low-diversity clusters in *BL* subsp. *infantis*, including the commercial probiotic strains BT1 (GCF_001281305.1, CM_MTB cluster) and EK3 (GCF_000730125.1, cMD3 cluster), as well as one porcine stool isolate (GCF_009696495.1, strain WCA-178-WT-4B) near the supplemented strain (**Figure 4A**). Conversely, multiple reference genomes (including commercial strains SC95, 139W423, W20-13, and BR3) grouped near low-diversity clusters in *B. breve*, but none were close to the supplemented strain (**Figure 4B**). Additional close matches were observed in *B. bifidum*, especially in a CM_MTB cluster (**Figure 4C**).

**Figure 4.**
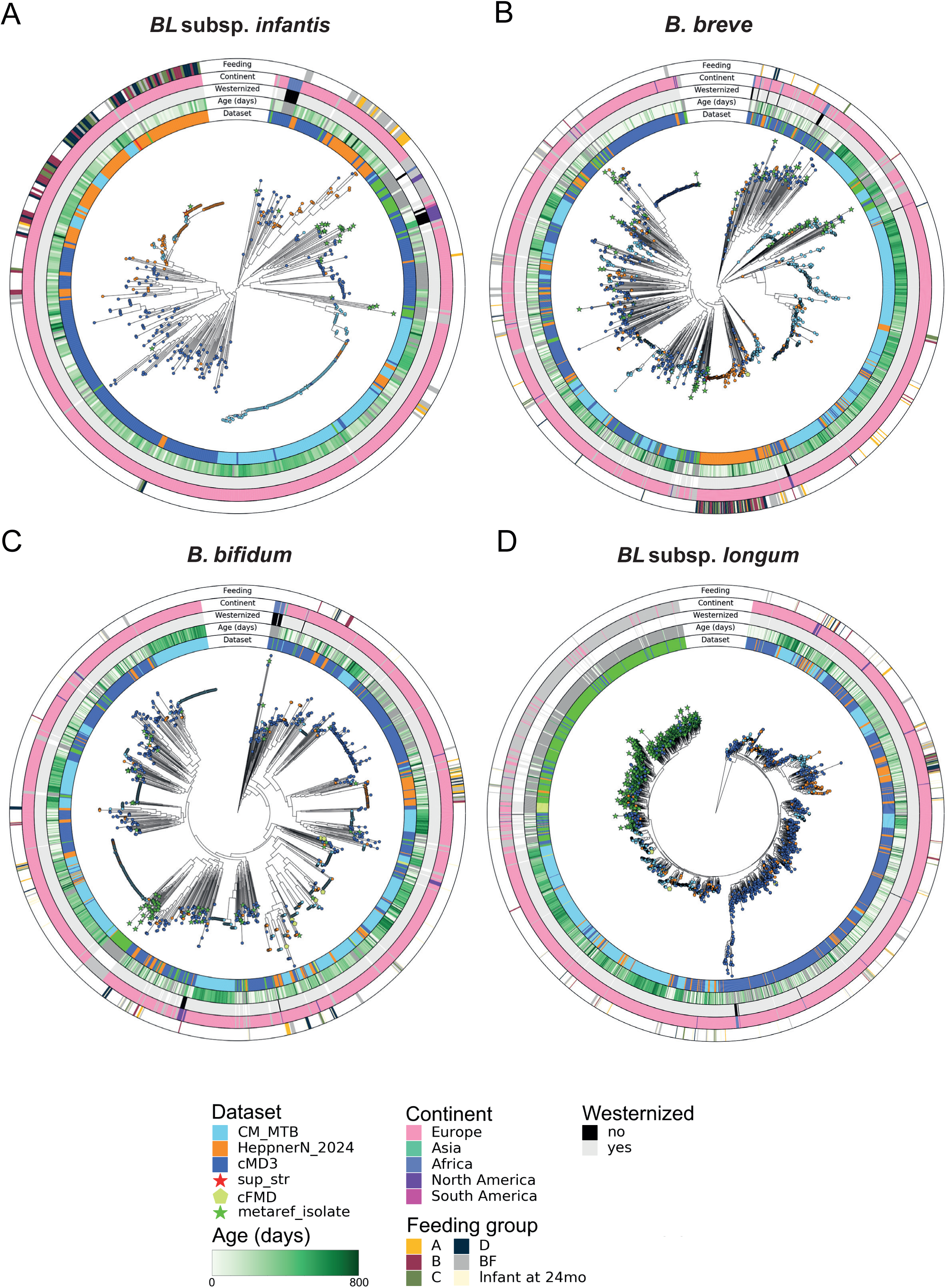
*Bifidobacterium* strain diversity in a global perspective. Phylogenies reconstructed from infant metagenomes (Infantibio-II/HeppnerN_2024, CM_MTB and cMD3 datasets, StrainPhlAn) containing each *Bifidobacterium* species, showing one dominant strain per sample. The phylogenies include (A) 543 samples for *BL* subsp. *infantis* (133 from Infantibio-II, 144 from CM_MTB, 237 from cMD3, 0 from cFMD, and 29 from MetaRefSGB) using 112 clade marker genes; (B) 1,438 samples for *B. breve* (207 from Infantibio-II, 566 from CM_MTB, 546 from cMD3, 3 from cFMD, and 116 from MetaRefSGB) using 199 clade marker genes; (C) 1,334 samples for *B. bifidum* (145 from Infantibio-II, 505 from CM_MTB, 594 from cMD3, 9 from cFMD, and 81 from MetaRefSGB) using 200 clade marker genes; and (D) 2,327 samples for *BL* subsp. *longum* (212 from Infantibio-II, 596 from CM_MTB, 988 from cMD3, 33 from cFMD, and 498 from MetaRefSGB) using 123 clade marker genes. The label “sup_str” (orange stars) indicates the supplemented *Bifidobacterium* reference strain, while “metaref_isolate” (green stars) represent other isolate reference genomes.

## Discussion

In this in-depth shotgun metagenomics analysis of gut microbiota from infants and paired parents from a randomized, controlled intervention trial, we demonstrate that pre- and probiotic exposure influences *Bifidobacterium* early life assembly at the species and strain levels. As previously shown^19^, formula- and breast-fed infants share characteristic features of microbiota assembly in the first year of life, developing towards the parental composition with age. Interestingly, we showed here that infant formula groups that were exposed to bifidobacteria (probiotics) supported an early life colonization of *BL* subsp. *infantis* at the first month of age, which was also observed with the GOS (prebiotics) supplemented group. This early dominance was absent in the breast-fed or placebo-formula groups – a pattern associated with the characteristic late or delayed colonization of *BL* subsp. *infantis* frequently observed in industrialized/westernized settings^22,39^. The rapid neonatal engraftment observed here appears to differ from reports describing limited colonization by many commercially available probiotics derived from high-income countries (HICs)^23^, although similar results have been reported for EVC001 strains in previous studies^40,41^. The absence of significant differences in species-level microbiome composition between feeding groups at later time points, together with the minor differences in Bray-Curtis dissimilarity at 24 months between groups, suggests that these effects are limited to early life and do not support stable long-term engraftment by these probiotic strains. These observations are consistent with strain-level findings reported by Bargheet et al. (2026), where the supplemented *BL* subsp. *infantis* strain initially dominated at 6 weeks of age, shortly after intervention ended, but was gradually replaced by endogenous strains by 6 months^42^.

Strain-level phylogenetic analysis of the two species related to the supplemented strains (*BL* subsp. *infantis* and *B. breve*) showed that infants exposed to the supplemented formula harboured a low-diversity cluster of prevalent strains, whereas breast-fed and placebo groups fell into a higher strain diversity cluster consistent with natural acquisition of endogenous strains. Notably, these patterns were recapitulated when integrating data from independent cohorts. Strikingly, a similar low-diversity *BL* subsp. *infantis* strain pattern was revealed for an Italian cohort (CM_MTB) independent of diet. This may suggest that formula supplementation with probiotics leads to a dominant colonization by the probiotic strain and antagonise strain diversity. This decrease of the intra-individual and inter-individuals dominant strain diversity would have an ecological consequence in the population, potentially hindering colonization by more diverse strains linked to breast-feeding.

In our study, the two *BL* subsp. *infantis* clusters (high and low diversity) were associated with distinct functional profiles, suggesting that this probiotic strain may possess specialised functional capacities that confer a temporary competitive advantage during early colonization, while naturally acquired strains maintain a broader functional repertoire typical of diverse microbial communities. The low-diversity highly-similar strain cluster – linked to probiotics exposure – contained more genes specialized in mobilization and recombination mechanisms. Strains belonging to this cluster were also enriched in antimicrobial activity mechanisms such as bacteriocin biosynthetic gene clusters (BGCs), including lantibiotic alpha and lanthionine synthetase C-like protein genes. *BL* subsp. *infantis* stands out among other bifidobacteria for carrying numerous BGCs, which likely provide a competitive advantage in colonising the infant gut by inhibiting rival microbes^43^. Conversely, a high-diversity strain cluster – related to breast-feeding and placebo - displayed a functional repertoire associated with transport and metabolism. Gene pathways related to carbohydrate degradation highly contributed by *Bifidobacterium* species, specifically *B. longum* and *B. breve* were enriched in formula-fed compared to breast-fed infants. Bifidobacterial-enriched gene families, including fructose-2,6-bisphosphate 2-phosphatase and alpha-glucosidase, which are associated with short-chain fatty acids (SCFA) production and carbohydrate degradation (including HMOs), respectively, were enriched in supplemented formula groups compared to placebo. SCFAs are critical immunological mediators that support T-cell regulation, suppress pro-inflammatory cytokine production, drive B-cell differentiation into IgA- and IgG-secreting plasma cells, and increase IgA coating of commensal bacteria^44^. Carbohydrate utilization capabilities vary across *Bifidobacterium* species and even among strains within the same species^23,45^. Consequently, pro- and prebiotics intervention studies should also consider strain-specific functional repertoires, not just species selection. Together, these findings support that infant formula supplementation does play a key role in bacterial strain diversity, vertical transmissibility and functional profiles.

In our cohort, breast-feeding was aligned with a higher maternal-to-offspring strain transmission compared with formula-feeding, as noted in similar studies^14^. We observed that this feeding-associated transmission pattern occurred for *BL* subsp. *longum* and *E. coli*, but not for *B. uniformis*. This suggests that this species may be transmitted by other routes independent of feeding mode. Vertical transmission has also been reported to be higher in vaginally delivered infants compared with those born by C-section^24^. Here, we don’t observe this phenomenon except for *BL* subsp. *longum* (data not shown). Interestingly, *BL* subsp. *infantis* differs from most other *Bifidobacterium* species in its mode of acquisition. Unlike many other *Bifidobacterium* species, which are commonly transmitted vertically from mother to infant, *BL* subsp. *infantis* is frequently acquired through horizontal transmission, such as nursery contacts^15^. Evidence suggests that its presence in the infant gut is strongly associated with the long-term breast-feeding history of a given population^46^. A distinct competitive advantage of *BL* subsp. *infantis* over other *Bifidobacterium* species lies in its specialized enzymatic machinery for HMO metabolism^47,48^. Among bifidobacteria, it has traditionally been the only species capable of utilizing all types of HMOs, yet its colonization in westernized populations occurs late and independently of maternal HMO composition^22,49–51^. A recent global genomic atlas^23^ demonstrated that *BL* subsp. *infantis* and *BL* subsp. *longum* represent ecologically distinct species and revealed pronounced biogeographic stratification of *BL* subsp. *infantis*, with substantially higher strain diversity in low- and middle-income countries (LMICs) compared to HICs. HIC-derived strains – including commercial probiotics (eg: EVC001) – are often clonal and miss key metabolic genes^23^. Although *BL* subsp. *infantis* is broadly associated with metabolism of breast milk-derived substrates such as HMOs, significant strain-level heterogeneity was observed, indicating that broad HMO utilization is not a universally conserved feature. LMIC-associated strains displays pronounced geo-specific metabolic adaptations to local diets, supporting the hypothesis of co-evolution with regional infant nutrition and weaning practices^23^.

IgA-microbe interactions, which play an important role in shaping microbial ecology and maintaining immune homeostasis, appeared to be largely determined by microbial taxonomy (genus- and species-specificity) rather than by dietary exposure in our cohort, as no significant differences in IgA coating were observed between feeding groups. Notably, *Veillonella* and *Escherichia* species consistently showed high IgA binding levels regardless of feeding mode and bacterial abundance. Interestingly, *B. bifidum*, but not *B. longum* or *B. breve*, was heavily targeted by IgA whenever it was detected, as seen previously^52^. Importantly, in our cohort, this trigger on IgA responses was not linked to dietary intervention, suggesting that IgA recognition is largely driven by taxonomic-specific properties rather than diet-associated microbial changes. Of note, IgA-coating of *Veillonella*, *Bifidobacterium*, *Bacteroides* and *Escherichia* genera was associated neither with total non-bound IgA in stool nor with the proportion of IgA-coated gut microbiota (data not shown). In another study performed with healthy adults, IgA binding was primarily driven by microbial gene content and was closely coordinated with other components of the immune system, rather than by microbial taxonomy alone^53^. In infancy, breastfeeding was associated with a higher level of IgA in the feces^52^. Human milk IgA binding to *BL* subsp. *infantis* was significantly higher in mothers from a low-allergy traditional farming community compared to urban mothers^52^. The absence of IgA-sorted metagenomes limited our ability to fully resolve strain- and gene-level characterization, although partial integration with whole-fecal metagenomic data helped strengthen our analyses. Another possible limitation is that from all the samples that were analysed for complete immunoglobulin phenotyping, only samples with a sufficient proportion of IgA-coated microbiota could be analyzed with IgA-Seq technology, thereby limiting the number of infants in each feeding or age group. Future studies using larger sample sizes and potentially IgA-sorted metagenomes could further help clarify the potential crosstalk between IgA surveillance, strain-level profiling and dietary intervention.

In the context of early-life microbiome development, *Bifidobacterium* dominance associated with supplement intake naturally reduced overall microbial diversity and richness. The reduced strain diversity observed in supplemented infants therefore most likely reflects successful competitive enrichment by the introduced probiotic strains rather than an actual microbiome dysfunction. Moreover, this is supported by the lack of elevated pathobionts or virulence markers. However, it still remains unclear whether the probiotic strains displace resident lineages, persist long-term, or influence other host functions.

## STAR Methods

### Resource availability

#### Lead contact

Further information and requests should be directed to and will be fulfilled by the lead contact Prof. Dr. Dirk Haller

### Materials availability

This study did not generate new unique reagents.

### Data and code availability

Sequencing data reported in this paper will be shared by the lead contact upon request.

This paper does not report original code at this time. Custom code used in this study will be deposited in a public repository prior to publication and will be made available upon request in the interim.

Any additional information required to reanalyze the data reported in this paper is available from the lead contact upon request.

### Method details

#### Sample collection

Fecal samples from the Infantibio-II cohort (including infants and their paired parents) were previously collected, processed and stored at -80°C^19^.

A longitudinal subset of samples (n = 60 infants; total number of infant samples = 297; total number of infant samples plus paired parental samples = 401):1-month (n = 59), 3-months (n = 59), 7-months (n = 60), 12-months (n = 59) and 24-months (n = 60) and paired parents (60 mothers and 44 fathers) collected when infants were 1-month old, was selected for shotgun metagenomics sequencing and downstream analyses (n = 401). Infants were delivered vaginally (n = 207) or by C-section (n = 90) (Table 1). Criteria used for samples selection was based on: complete longitudinal series across all time points, balanced number of samples between the formula groups and no antibiotics used in any infants. Here we compared a breast-fed (BF) reference group with four formula-fed intervention groups receiving: placebo (formula A), bifidobacteria (probiotics, formula B), galacto-oligosaccharides (GOS; prebiotics, formula C), or a synbiotic combination of both (formula D). “Feeding group” refers to the feeding received by the infant at the considered time-point, or over the study course for 24-months samples. Formula groups across the first-year infants (n = 237) consisted of: formula A (n = 30), formula B (n = 34), formula C (n = 28) and formula D (n = 35). Breast-fed samples (n = 110) were used as control (Table 1).

**Table 1.** Study design including groups by age and delivery mode.

| INFANTIBIO-II INFANT-PARENTAL METAGENOMES |  |  |  |
| --- | --- | --- | --- |
| Infant sampling time point or parent | Born via vaginal route (n = 207) | Born via C-section (n = 90) | Total (n = 401) |
| 1-month | 41 | 18 | 59 |
| 3-months | 41 | 18 | 59 |
| 7-months | 42 | 18 | 60 |
| 12-months | 41 | 18 | 59 |
| 24-months | 42 | 18 | 60 |
| Mothers | - | - | 60 |
| Fathers | - | - | 44 |

**Table 2.** Study design including groups by feeding exposure.

| Feeding group | Sampling time point |  |  |  |  |
| --- | --- | --- | --- | --- | --- |
|  | 1-month<br>(n = 59) | 3-months<br>(n = 59) | 7-months<br>(n = 60) | 12-months<br>(n = 59) | Follow-up at 24-months<br>(n = 60) |
| Formula A | 5 | 6 | 9 | 10 | 10 |
| Formula B | 7 | 7 | 10 | 10 | 10 |
| Formula C | 5 | 5 | 8 | 10 | 10 |
| Formula D | 7 | 8 | 10 | 10 | 10 |
| Breast-fed | 35 | 33 | 23 | 19 | 20 |

#### Genomic DNA isolation and metagenomic sequencing

The bacterial lysis and genomic DNA isolation were carried in accordance with a modified protocol by Godon *et al*. (1997)^54^. In brief, 2 mL screw-cap tubes were filled with 400 mg of 0.1 mm zirconium/glass pellets (Roth, Ref. Nr.: N033.1) and autoclaved at 121 °C for 20 min for bacterial lysis. Chemical lysis was performed by filling the tubes in sterile conditions with 250 μL 4 M Guanidinethiocyanat (Sigma, Ref. 11 Nr.: 593840, diluted in 0.1 M Tris) and 500 μL 5% N-lauroylsarcosine (Sigma, Ref. Nr.: 137166, dissolved in DPBS). Remaining microbial cells were lysed using a FastPrep-24 fitted with a CoolPrep adapter filled with dry ice. gDNA was purified using a NucleoSpin gDNA Clean-up kit (Macherey-Nagel, Ref. Nr.: 740230.250).

DNA was randomly sheared and paired-end libraries were prepared by ligating Illumina-compatible adapters to the fragmented DNA. Library quantification was carried out using Qubit and quality assessment was conducted via BioAnalyzer to ensure optimal library preparation. Sequencing was performed on an Illumina NovaSeq 6000 platform (paired-end 150 bp), with a target output of 12 Gb of high-quality reads per demultiplexed sample.

#### lmmunoglobulin phenotyping, lgA sorting and 16S rRNA sequencing

Fecal samples from 129 mother-infant dyads (n = 597 total samples) were selected for immunoglobulin characterization. IgA sorting was performed on 135 samples from 43 dyads, including 81 samples previously analyzed by whole-genome deep shotgun metagenomics (see above).

##### Microbiota purification

Briefly, stool samples were resuspended in cold 1xPBS. Large debris were pelleted by centrifugation at 300rpm for 15min. Supernatant was centrifuged at 3500rpm for 15min. Supernatant was removed and pellet resuspended in 1xPBS containing 10% glycerol. Samples were stored at -80°C for downstream analysis.

##### lmmunoglobulin phenotyping

Approximately 10^7^ microbes were stained with F(ab’)2 goat anti-human IgA, IgG and IgM antibodies as well as Fc fragment (conjugated to FITC, DL405, AF594 and AF647, respectively; Jackson-Immunoresearch, France). Single-cell fluorescent intensity was measured by flow cytometry (Cytoflex S, Beckman Coulter) and analysed with FlowJO (BD, France) and FunkyCells (www.funkycells.com) softwares.

##### lgA sorting

Samples with proportions of IgA-coated microbiota exceeding 10% were sorted. Briefly, 150 µL of purified stool samples were incubated with FITC-conjugated F(ab’)2 goat anti-human IgA antibody (Jackson ImmunoResearch, France) for 20 min at room temperature in the dark. Samples were washed with 1 mL of 1xPBS and centrifuged at 9600 rpm for 10 min at 4°C. The pellets were resuspended in 480 µL of 1xPBS and incubated with 20 µL of anti-FITC magnetic microbeads (Miltenyi Biotec, France) for 20 minutes at room temperature in the dark. Magnetic enrichment was performed using a MultiMACS Cell 24 system (Miltenyi Biotech, France) according to the manufacturer instructions. Positive and negative fractions were centrifuged at 14,000 rpm for 10 minutes and their pellets were stored at -80°C. A second column pass was performed for negative fractions showing IgA labeling.

##### DNA extraction and 16S rRNA gene sequencing

DNA extractions were performed using the ZymoBIOMICS 96 MagBead DNA Kit according to the manufacturer instructions. Samples were shaken (10,000 rpm) three times for 1 min using the Precellys Evolution tissue homogenizer (Bertin). DNA was resuspended in 50 µL of DNase/RNase free water and stored at −20°C until amplification.

The hypervariable V3-V4 regions of the 16S rRNA gene were amplified from extracted DNA using a semi-nested PCR approach^55^. In the first PCR, a 720 bp fragment (V3-V6 regions) was amplified using the primers bakt_341F (5’- CCTACGGGNGGCWGCAG -3’) and 1061R (5’-CRRCACGAGCTGACGAC -3). The amplification was carried out over 10 cycles with the following conditions: denaturation at 94 °C, hybridization at 55 °C, and elongation at 72 °C. The second amplification targeting the V3–V4 regions was performed using bakt_341F_ICM (5’-AAGACTCGGCAGCATCTCCATCCTACGGGNGGCWGCAG-3’) and bakt_805R_ICM (5’-GCGATCGTCACTGTTCTCCAATCTGACTACHVGGGTATCTAATCC-3’) primers with a denaturation temperature of 94°C, hybridization at 58°C, and elongation at 72°C. Sequencing was done with Illumina NextSEQ 2x300bp paired-end sequencing technology (ICM, Paris, France).

##### Bioinformatics and lgA-Seq scores

For each sorted sample, three fractions were obtained: one IgA-positive fraction and two IgA-negative fractions (the negative fraction was subdivided into 90% and 10% aliquots). The use of two negative fractions was intended to quantify downstream experimental variability independent of biological differences. To model technical dispersion, ASV abundances from the two negative fractions (neg1 and neg2) were compared. For each ASV, log₂(neg1/neg2) was plotted against log₁₀(neg1 x neg2). Because technical variation depends on signal intensity, sliding windows along the log₁₀(neg1 x neg2) axis were defined, each containing 20 ASVs. Within each window, the distribution of log₂ ratios was centered and scaled to generate a standardized Gaussian distribution (Z-transformation), thereby estimating the intensity-dependent experimental variation.

Using this intensity adjusted null distribution derived from the comparison of the two biologically identical negative fractions, a standardized IgA-Seq Z-score was calculated for each ASV when comparing the IgA-positive and IgA-negative fractions. ASVs with Z-scores exceeding ±1.645 (α = 0.10) were considered significantly differentially abundant: Z > +1.645 indicated enrichment in the IgA-positive fraction, whereas Z < -1.645 indicated enrichment in the IgA-negative fraction.

For the circos plot representation, relative abundances were normalized using the min-max method. IgA scores were transformed using the formula: 0.5 + (medianscore / 2xmax(min_val, max_val)) to create a score centered around 0.5 and restrained to the interval between 0 and 1.

#### Metagenomic analysis and assembly

Quality control (QC), including adapter trimming, low-quality filtering and removal of host-derived reads was conducted using Trim Galore v 0.6.10^56^ and KneadData v 0.10.0^57^. For adapter removal and low-quality filtering, Trim Galore was run using the parameters “--paired --phred33 --quality 0 --stringency 5 --length 10”. The subsequently applied tool KneadData includes also trimming with the built-in read trimmer Trimmomatic but is mainly applied for removal of human contaminant reads. The parameters “--trimmomatic-options ’HEADCROP:15 SLIDINGWINDOW:4:15 MINLEN:50’“ and bowtie2 with the human reference genome hg38 were used.

On the clean reads, reference-based taxonomic profiling was performed using MetaPhlAn4 v 4.1.1^38^ with a tailored reference database to accurately differentiate between *B. longum* subspecies (markers added to the mpa_vOct22_CHOCOPhlAnSGB_202212 database^22^). To enrich this data, we also considered 651 healthy infant (< 2.5 years old) stool samples from the microTOUCH-baby (CM_MTB) cohort^15^, 2,151 healthy stool samples from the cMD3 database (curatedMetagenomicData, v. 3.18^36^), filtered by age (< 2.5 years old) and without antibiotic intake, 76 food metagenomic samples from the cFMD database (curatedFoodMetagenomicData, v 1.2.1^37^) that had at least one of the four *Bifidobacterium* species/subspecies of interest present in their taxonomic profile. We also queried MetaRefSGB^37,38^, a microbial genomic database containing >219,000 isolate genomes and >1,300,000 metagenome-assembled genomes (MAGs) as of vJan25, for isolate genomes or MAGs from food sources, and included a total of 724 isolate reference genomes (*BL* subsp. *Infantis* n=29 ; *BL* subsp. *longum* n=498 ; *B. breve* n=116 ; *B. bifidum* n=81) in the trees as references. These integrated datasets were processed using the same custom marker gene database; the resulting consensus marker sequences together with the clade-specific marker genes were used together with our data for strain-level analysis and building phylogenetic trees. Strain-level analysis related to single-nucleotide polymorphisms (SNPs) variation was carried out using StrainPhlAn4 v 4.1.1^58^ (default parameters) which leverages species-specific marker genes for strain identification and strain tracking across samples. The StrainPhlAn runs produced RAxML^59^ maximum likelihood .tre files in Newick format, which were visualized and rooted at midpoint using iTOL^60^. The new trees were annotated using GraPhlAn^61^ with custom annotations and decorated with rings representing the sample’s dataset, age (when available), continent, westernization, and feeding mode. Evolutionary distances were obtained using the EMBOSS distmat package v 6.6.0.0 with the Kimura correction. Functional profiling was performed using HUMAnN3 v 3.6^57^ in combination with results from MetaPhlAn3 v 3.1.0^57^. The UniRef90 protein database was used for gene family mapping, and gene pathways and gene family abundance units were normalized to Copies Per Million (CPM). Results from individual samples were joined. Gene families were additionally regrouped and renamed according to MetaCyc reactions^62^.

De-novo assembly of clean reads into contigs was undertaken using MEGAHIT v 1.2.9^63^ with default parameters and protein-coding genes were predicted using Prodigal v 2.6.3^64^, including those from whole genome sequencing data from the *BL* subsp. *infantis* supplemented strain^65^. For Prodigal, option “-p meta” was chosen to use the optimal mode for metagenomic data. After filtering incomplete genes with the “partial==00” flag, cd-hit-est v 4.8.1^66^ was used to cluster all genes based on at least 95% identity and 90% of coverage using the parameters “-aS 0.9 -aL 0.9 -c 0.95 -d 0”. The representative genes for all gene families were translated into protein sequences and re-clustered using cd-hit v 4.8.1 by 90% identity and 80% coverage of amino acid sequence alignments (parameters “-d 100 -c 0.9 -aL 0.8 -aS 0.8 -G 0”).

The representative genes were clustered into metagenomic species pangenomes (MSPs) based on co-abundance of these representative genes in the samples using MSPminer^67^ with its default parameters. The MSPs were then taxonomically annotated using GTDB-Tk v 2.3.2^68^ for species-level annotations. Since species-level annotations were not possible for many MSPs, missing taxonomic annotations were supplemented taking into account the reference-based results from MetaPhlAn 4^69^. Annotations of protein functions based on functional domains were assessed using InterProScan v 5.55_88.0^70^. For each representative protein from the non-redundant catalog, Pfam domain annotations were predicted (“-appl Pfam”).

Metagenomic data were processed using an in-house pipeline implemented in Snakemake v 7.26.0^71^, applying the tool versions and parameters specified above. The overall workflow and parametrization are highly consistent with our recently published MetaGear implementation in Nextflow ^72^, available at: https://github.com/schirmer-lab/metagear-pipeline.

#### Statistical analysis and data visualization

Alpha diversity parameters and dissimilarity indices were obtained using the vegan package v 2.7-2^73^ from R^74^. For beta diversity, Bray-Curtis was used as dissimilarity metric and PERMANOVA was performed with the adonis2 function from the vegan package (infant IDs as “blocks” for permutations to account for non-independence of samples) to test the interaction of feeding and age, adjusting for sex, delivery mode (“vaginal” or “c-section”) and maternal antibiotics use (“yes” or “no”), using ‘by = "margin"’. Pairwise PERMANOVA^75^ was then applied to compare feeding (with all time points together or within each time point) or age (with all time points together) groups with BH correction^33^. Homogeneity of multivariate dispersions among feeding or age groups was assessed using betadisper (vegan package). Kruskal Wallis^34^ (stats package) and posthoc Dunn (dunn.test package^76^) were used to assess the effect between feeding exposure and *Bifidobacterium* species relative abundances. Benjamini-Hochberg (BH) correction for p-values and prevalence filtering (n=4) were always performed unless stated otherwise. Downstream analyses and visualization were utilized using in-house R v 4.4.0^74^ scripts with dplyr v 1.1.4^77^ and ggplot2 v 3.5.2^78^ and in-house Python v 3.12.1 scripts with numpy v 1.24.3^79^, matplotlib v 3.7.2^80^, seaborn v 0.11.2^81^, pandas v 1.5.3^82^, scipy v 1.11.1^83^ and statannotations v 0.6.0^84^. Seaborn “deep” color palette was used for the *Bifidobacterium* species boxplots.

Low-diversity cluster 2 for *BL* subsp. *infantis* was defined by strain identity to the reference probiotic strain derived from Kimura 2-parameter-corrected distances (>96.02% for Infantibio-II phylogeny; >96.84% for the combined phylogeny of Infantibio-II and BMB samples). Corresponding thresholds for *B. breve* were >97.28% and >97.55%, respectively. Hypergeometric distribution (R stats package (v 4.4.0)) with BH correction was conducted to determine over-representation of feeding groups in the *BL* subsp. *infantis* and *B. breve* respective clusters. Non-parametric Mann-Whitney^85^ (or Kruskal-Wallis^34^ if stated otherwise) were applied for statistical testing of parents-to-offspring evolutionary distances between metadata groups (relationship, feeding or sampling time point).

Multivariate analyses was carried out using MaAsLin2 v 1.18.0^86^ with the parameters: min_abundance = 10 (for gene families) or 1000 (for gene pathways); transform = “LOG”; normalization = NONE; analysis_method = “LM”; correction = “BH”; max_significance = 0.05 (for gene families) and 0.25 (for gene pathways); with the fixed effects (covariates) for the Placebo (formula group A) vs. Supplementation (groups B, C, and D) gene families: probiotics/prebiotics supplementation (“no” as reference), feeding (“bf” as reference), antibiotics in mother (“no” as reference), delivery mode (“vaginal” as reference) and sex (“female” as reference); fixed effects for the Cluster 1 vs. Cluster 2 gene families: Cluster (“1” as reference), feeding (“bf” as reference), antibiotics in mother (“no” as reference), delivery mode (“vaginal” as reference) and sex (“female” as reference); fixed effects for gene pathways: feeding (“bf” as reference), antibiotics in mother (“no” as reference), delivery mode (“vaginal” as reference) and sex (“female” as reference); and subject IDs as random effects. Additionally, 10% of prevalence was applied before this run for the clustering. Packages ggrepel v 0.9.6^87^ and ggtext v 0.1.2^88^ were utilized for HUMAnN3 volcano plots among others previously stated. Stratified HUMAnN3 plots were generated using the command humann_barplot.

## Acknowledgements

This study was funded by the Joint Program Initiative of the European Union (project name EcoBiotic) and the German Ministry of Education and Research (BMBF; FKZ 01EA2207).

The Technical University of Munich provided technical support through the Core Facility Microbiome of the ZIEL Institute for Food & Health (Klaus Neuhaus, Aritra Mahapatra, Lukas Mix, Angela Sachsenhauser and Caroline Ziegler) for shotgun metagenomics sequencing and through DaiSyBio (Data Science in Systems Biology, Markus List) for server space.

## Author contributions

D.H initiated and coordinated the study, reviewed all data and secured funding, M.S supervised the shotgun metagenomics analyses and provided additional server space and computational power, M.L supervised the immunoglobulin phenotyping and 16S rRNA amplicon sequencing of the IgA sorted data, E.D performed the IgA-sorted data analyses, M.C.C provided the *Bifidobacterium* strains, T.H and L.L performed the antibody measurements, R.S carried out the global strain diversity analyses, N.S and V.H supervised the implementation and analyses of the global strain diversity workflow, M.V.M was responsible for performing shotgun metagenomics analyses of the whole fecal samples, including statistical analysis, data generation and data visualization, M.V.M and S.W carried out the execution of the metagenomics pipeline tools, M.V.M, A.E, H.O, M.S and D.H wrote the manuscript.

## Supplemental information

**Figure S1.**
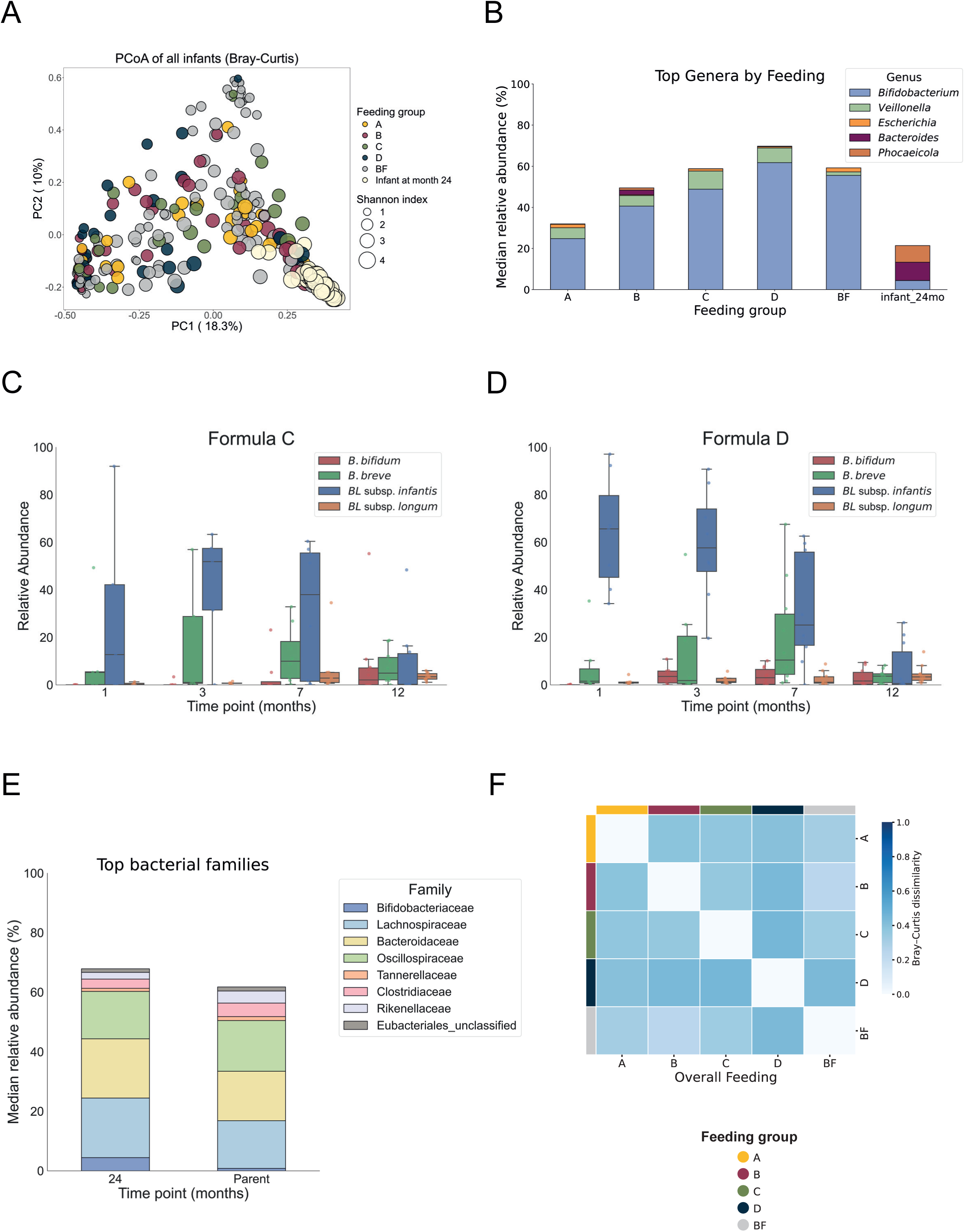
(A) PCoA plot of beta-diversity (Bray-Curtis) of all infant samples (n= 297) grouped according to feeding modality by color with Shannon diversity (alpha-diversity) shown by size. (B) Distribution of the top genera of all infants by median relative abundance (%) with each bar representing one feeding group. First-year infants (n = 237) are stratified by feeding group (formula groups A-D and breast-fed/BF), while the samples at month 24 are represented in a separate bar. (C) Top *Bifidobacterium* species by median relative abundance (%) for formula C-fed infants with each color representing each species. (D) Top *Bifidobacterium* species by median relative abundance (%) for formula D-fed infants with each color representing each species. (E) Taxonomic distribution at the family-level of the 24-months infant samples and the parental samples by color. Bacterial families with <1% relative abundance were not considered. (F) Heatmap of 24-months infant samples with Bray-Curtis dissimilarity (based on median taxonomic abundances within each feeding group) between each overall feeding group. Overall feeding refers to the feeding groups to which infants were assigned during the first year of life (the intervention period). The long-term effect of this early intervention is evaluated here at 24 months of age.

**Figure S2.**
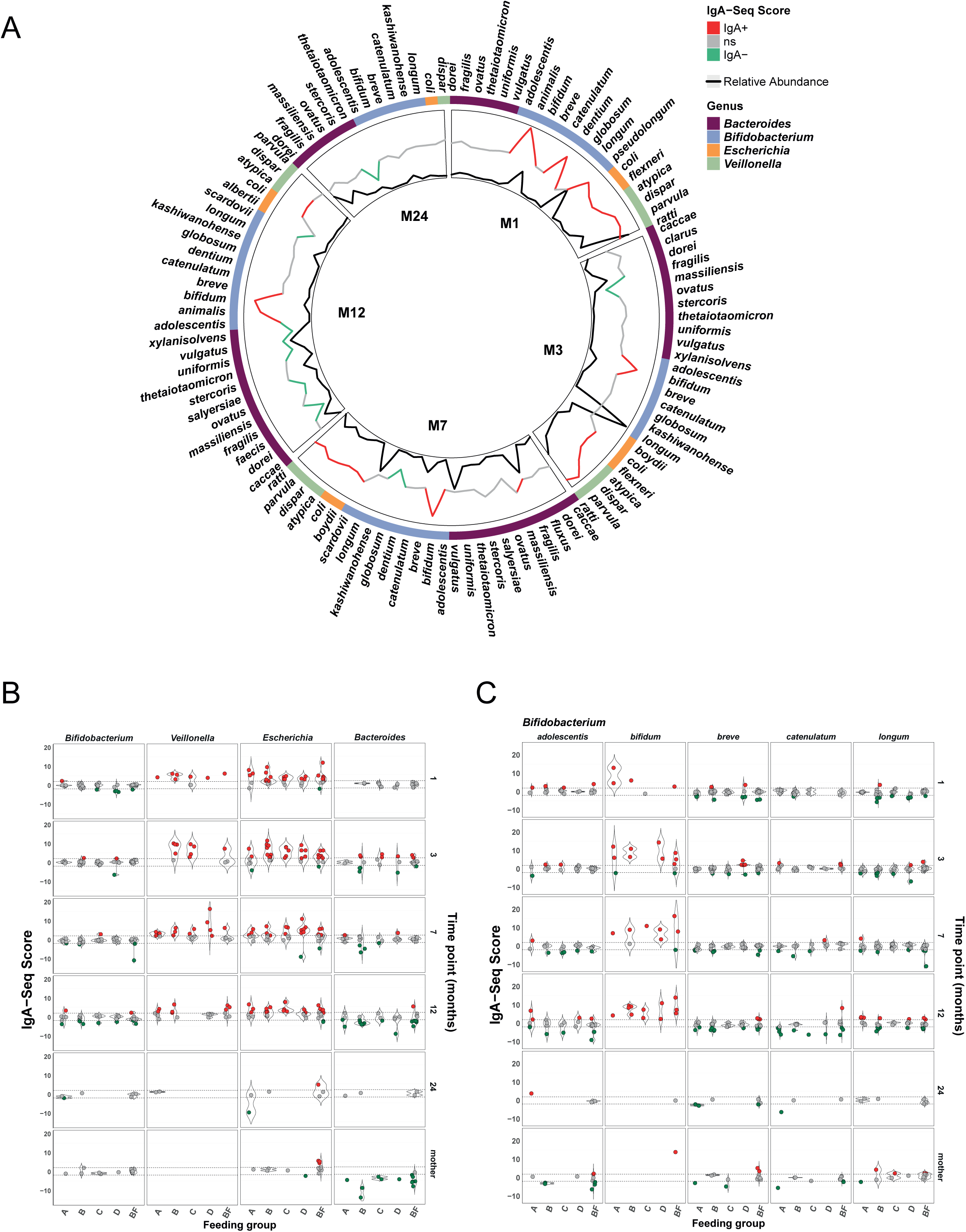
(A) Circos plot of all infant samples of 16S rRNA sequenced IgA-sorted data displaying IgA-binding scores for all bacterial taxa at species-level (outer layer) and bacterial relative abundances (inner layer). (B) Median IgA-binding scores for the dominant bacterial taxa (genus-level) in infants sorted by sampling time point (y-axis) and distributed by feeding groups (x-axis). (C) Median IgA-binding scores for the *Bifidobacterium* species in infants sorted by sampling time point (y-axis) and distributed by feeding groups (x-axis).

**Figure S3.**
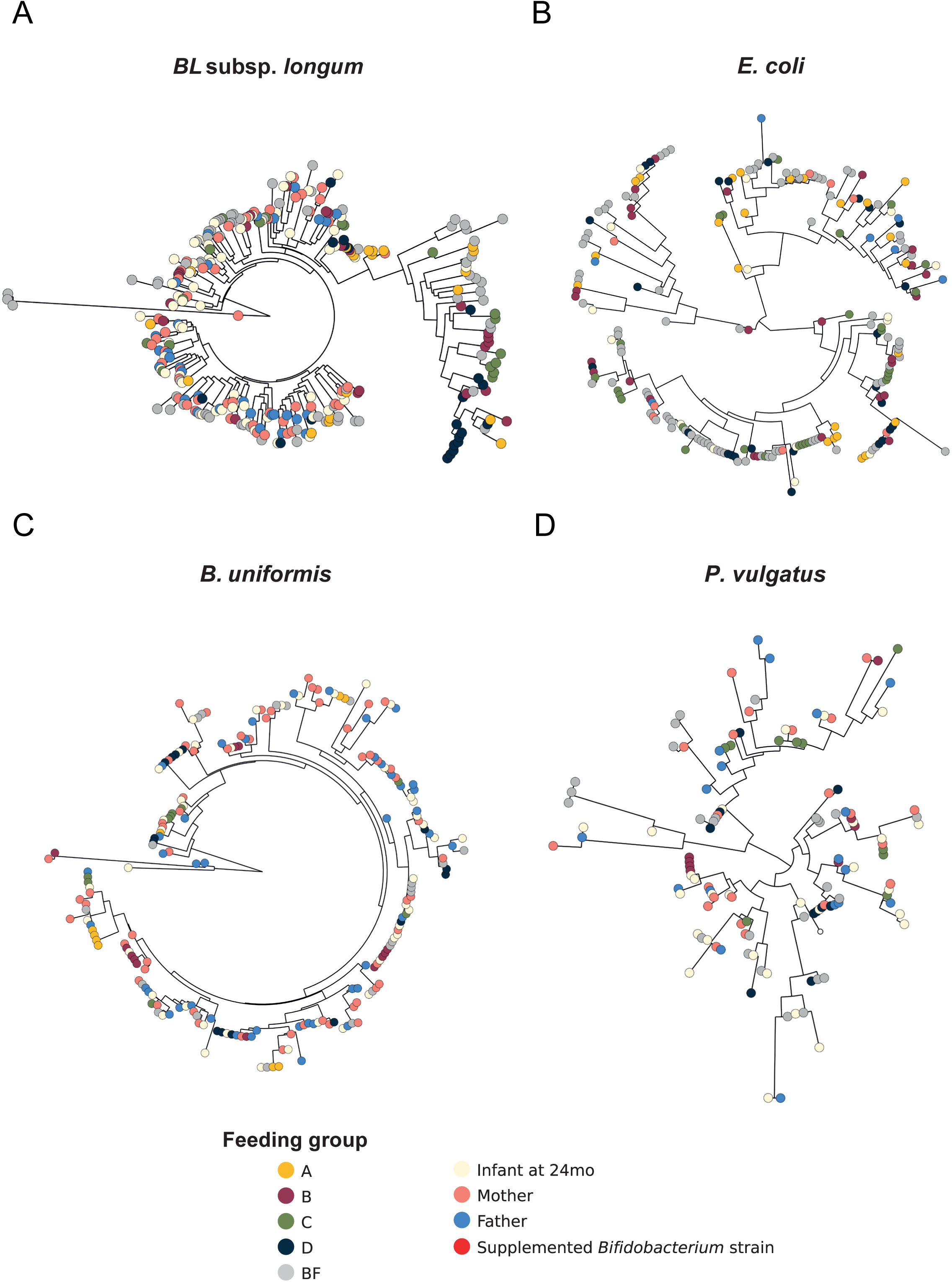
StrainPhlAn phylogenies reconstructed from metagenomes of infants and their paired parents (n = 401) showing one dominant strain per sample. Phylogenies were constructed for (A) 293 samples for *BL* subsp. *longum* using 128 clade marker genes, (B) 218 samples for *E. coli* using 128 clade marker genes, (C) 209 samples for *B. uniformis* using 136 clade marker genes, and (D) 123 samples for *P. vulgatus* using 78 clade marker genes. Feeding mode is indicated by color.

**Figure S4.**
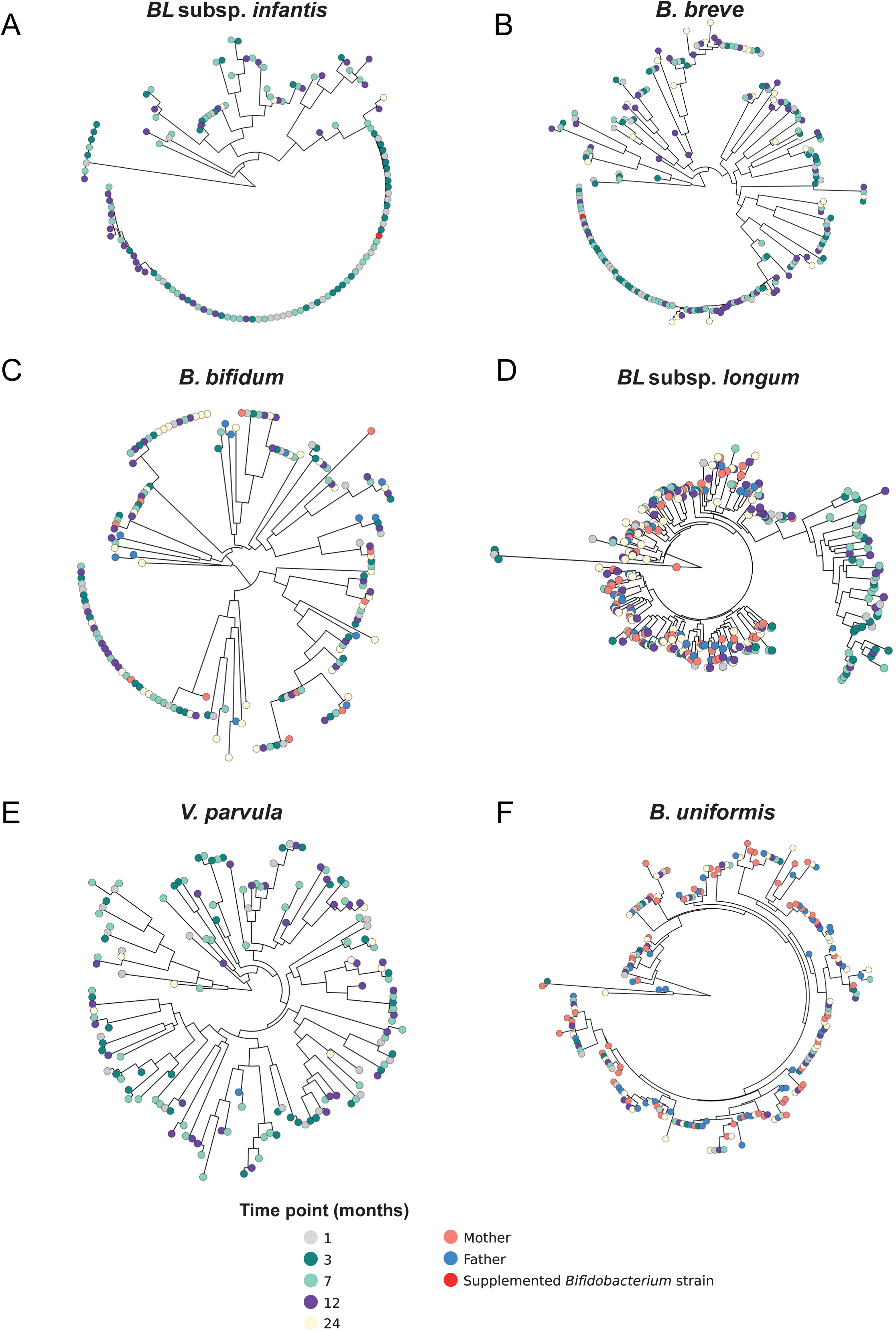
StrainPhlAn phylogenies reconstructed from metagenomes of infants and their paired parents (n = 401), showing one dominant strain per sample. Phylogenies were constructed for (A) 133 samples for *BL* subsp. *infantis* using 119 clade marker genes, (B) 207 samples for *B. breve* using 200 clade marker genes, (C) 168 samples for *B. bifidum* using 200 clade marker genes, (D) 293 samples for *BL* subsp. *longum* using 128 clade marker genes, (E) 148 samples for *V. parvula* using 200 clade marker genes, and (F) 209 samples for *B. uniformis* using 136 clade marker genes. Time point is indicated by color. Supplemented *Bifidobacterium* strains (*BL* subsp. *infantis* and *B. breve*) are highlighted in red as reference strains.

**Figure S5.**
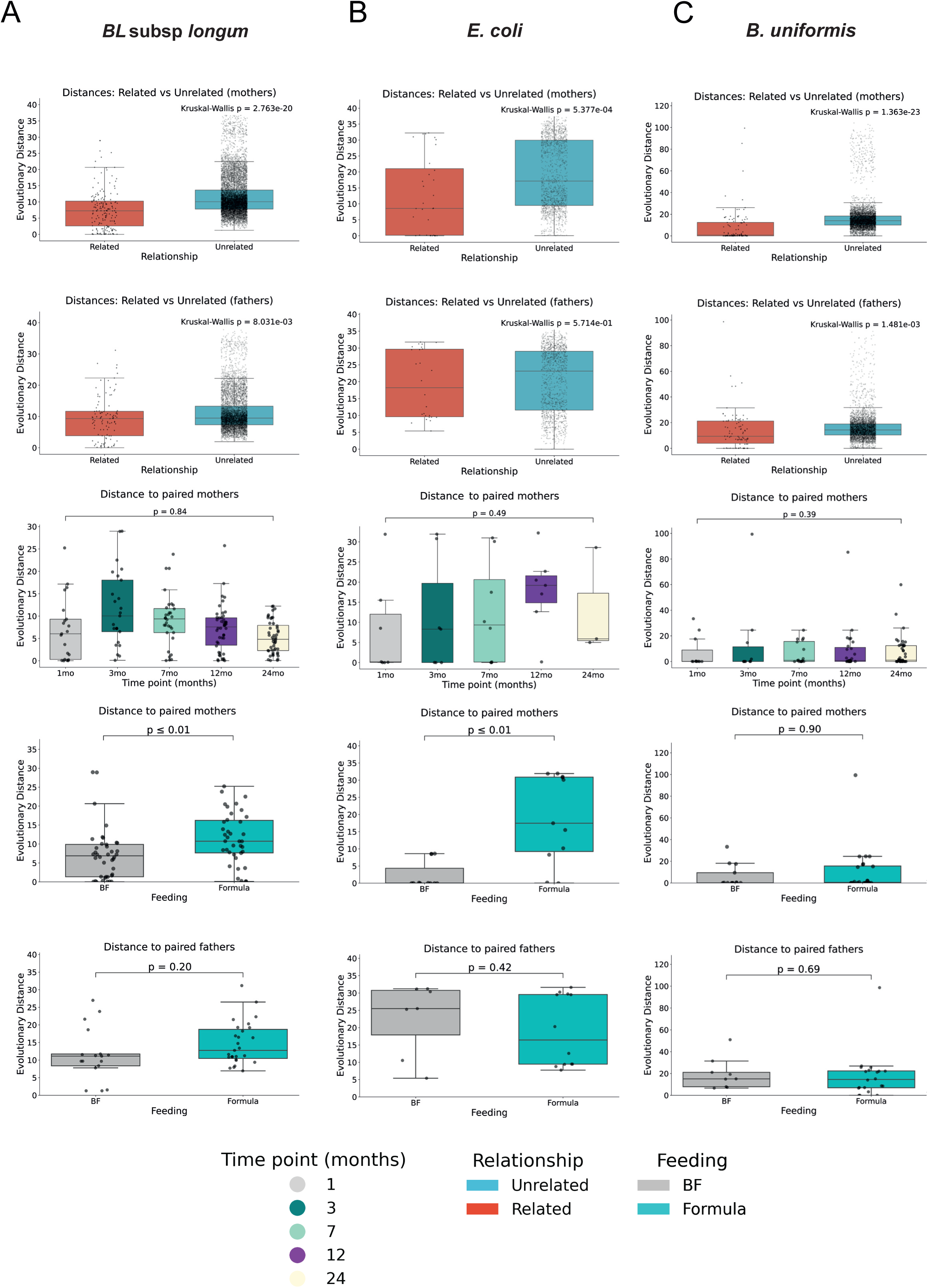
StrainPhlAn evolutionary distances (Kimura-corrected) from infants to parents are shown for (A) BL subsp. longum, (B) E. coli, and (C) B. uniformis. Top panels display distances between infants and mothers or fathers (related vs. unrelated), with significance assessed using the Kruskal-Wallis test. Distances to paired mothers are stratified by sampling time point and highlighted by color. The lower panels show evolutionary distances of 1-, 3-, and 7-month samples to paired mothers and fathers stratified by dietary intervention (breast-fed/BF vs formula). Statistical comparisons for these subplots were performed using the Mann-Whitney test (with Benjamini-Hochberg correction).

## References

1. Backhed, F., Roswall, J., Peng, Y., Feng, Q., Jia, H., Kovatcheva-Datchary, P., Li, Y., Xia, Y., Xie, H., Zhong, H., et al. (2015). Dynamics and Stabilization of the Human Gut Microbiome during the First Year of Life. Cell Host Microbe 17, 690–703. 10.1016/j.chom.2015.04.004.

2. Biasucci, G., Rubini, M., Riboni, S., Morelli, L., Bessi, E., and Retetangos, C. (2010). Mode of delivery affects the bacterial community in the newborn gut. Early Hum. Dev. 86, 13–15. 10.1016/j.earlhumdev.2010.01.004.

3. Dominguez-Bello, M.G., Costello, E.K., Contreras, M., Magris, M., Hidalgo, G., Fierer, N., and Knight, R. (2010). Delivery mode shapes the acquisition and structure of the initial microbiota across multiple body habitats in newborns. Proc. Natl. Acad. Sci. U. S. A. 107, 11971–11975. 10.1073/pnas.1002601107.

4. Intze, E., Schaubeck, M., Pourjam, M., Neuhaus, K., Lagkouvardos, I., Hitch, T.C.A., and Clavel, T. (2025). The infant microbiota hopscotches between community states toward maturation—longitudinal stool parameters and microbiota development in a cohort of European toddlers. ISME Commun. 5. 10.1093/ismeco/ycaf016.

5. Stewart, C.J., Ajami, N.J., O’Brien, J.L., Hutchinson, D.S., Smith, D.P., Wong, M.C., Ross, M.C., Lloyd, R.E., Doddapaneni, H.V., Metcalf, G.A., et al. (2018). Temporal development of the gut microbiome in early childhood from the TEDDY study. Nature 562, 583. 10.1038/s41586-018-0617-x.

6. Yassour, M., Vatanen, T., Siljander, H., Hamalainen, A.M., Harkonen, T., Ryhanen, S.J., Franzosa, E.A., Vlamakis, H., Huttenhower, C., Gevers, D., et al. (2016). Natural history of the infant gut microbiome and impact of antibiotic treatment on bacterial strain diversity and stability. Sci. Transl. Med. 8. 10.1126/scitranslmed.aad0917.

7. Bergstrom, A., Skov, T.H., Bahl, M.I., Roager, H.M., Christensen, L.B., Ejlerskov, K.T., Mølgaard, C., Michaelsen, K.F., and Licht, T.R. (2014). Establishment of intestinal microbiota during early life: a longitudinal, explorative study of a large cohort of Danish infants. Appl. Environ. Microbiol. 80, 2889–2900. 10.1128/AEM.00342-14.

8. Ojima, M.N., Jiang, L., Arzamasov, A.A., Yoshida, K., Odamaki, T., Xiao, J., Nakajima, A., Kitaoka, M., Hirose, J., Urashima, T., et al. (2022). Priority effects shape the structure of infant-type Bifidobacterium communities on human milk oligosaccharides. ISME J. 16, 2265–2279. 10.1038/s41396-022-01270-3.

9. Shao, Y., Garcia-Mauriiio, C., Clare, S., Dawson, N.J.R., Mu, A., Adoum, A., Harcourt, K., Liu, J., Browne, H.P., Stares, M.D., et al. (2024). Primary succession of Bifidobacteria drives pathogen resistance in neonatal microbiota assembly. Nat. Microbiol. 2024 910 *9*, 2570–2582. 10.1038/s41564-024-01804-9.

10. Sprockett, D., Fukami, T., and Relman, D.A. (2018). Role of priority effects in the early-life assembly of the gut microbiota. Nat. Rev. Gastroenterol. Hepatol. 15, 197–205. 10.1038/nrgastro.2017.173.

11. Feehily, C., O’Neill, I.J., Walsh, C.J., Moore, R.L., Killeen, S.L., Geraghty, A.A., Lawton, E.M., Byrne, D., Sanchez-Gallardo, R., Nori, S.R.C., et al. (2023). Detailed mapping of Bifidobacterium strain transmission from mother to infant via a dual culture-based and metagenomic approach. Nat. Commun. 14. 10.1038/s41467-023-38694-0.

12. Sawhney, S.S., Thanert, R., Thanert, A., Hall-Moore, C., Ndao, I.M., Mahmud, B., Warner, B.B., Tarr, P.I., and Dantas, G. (2025). Gut microbiome evolution from infancy to 8 years of age. Nat. Med. 31, 2004. 10.1038/s41591-025-03610-0.

13. Vatanen, T., Jabbar, K.S., Ruohtula, T., Honkanen, J., Avila-Pacheco, J., Siljander, H., Stražar, M., Oikarinen, S., Hyöty, H., Ilonen, J., et al. (2022). Mobile genetic elements from the maternal microbiome shape infant gut microbial assembly and metabolism. Cell 185, 4921–4936.e15. 10.1016/j.cell.2022.11.023.

14. Yassour, M., Jason, E., Hogstrom, L.J., Arthur, T.D., Tripathi, S., Siljander, H., Selvenius, J., Oikarinen, S., Hyoty, H., Virtanen, S.M., et al. (2018). Strain-Level Analysis of Mother-to-Child Bacterial Transmission during the First Few Months of Life. Cell Host Microbe 24, 146–154.e4. 10.1016/j.chom.2018.06.007.

15. Ricci, L., Heidrich, V., Punčochář, M., Armanini, F., Ciciani, M., Nabinejad, A., Fazaeli, F., Piperni, E., Servais, C., Pinto, F., et al. (2026). Baby-to-baby strain transmission shapes the developing gut microbiome. Nat. 2026, 1–10. 10.1038/s41586-025-09983-z.

16. Adjele, J.J.B., Devi, P., Kumari, P., Yadav, A., Tchuenchieu Kamgain, A.D., Mouafo, H.T., Medoua, G.N., Essia, J.J.N., Chauhan, N.S., and Pandey, R. (2024). Exploring the influence of age and diet on gut microbiota development in children during the first 5 years: a study from Yaoundé, Cameroon. Front. Microbiol. 15, 1512111. 10.3389/fmicb.2024.1512111.

17. Fouhy, F., Watkins, C., Hill, C.J., O’Shea, C.A., Nagle, B., Dempsey, E.M., O’Toole, P.W., Ross, R.P., Ryan, C.A., and Stanton, C. (2019). Perinatal factors affect the gut microbiota up to four years after birth. Nat. Commun. 2019 101 10, 1517-. 10.1038/s41467-019-09252-4.

18. Ma, J., Li, Z., Zhang, W., Zhang, C., Zhang, Y., Mei, H., Zhuo, N., Wang, H., Wang, L., and Wu, D. (2020). Comparison of gut microbiota in exclusively breast-fed and formula-fed babies: a study of 91 term infants. Sci. Reports 2020 101 *10*, 15792-. 10.1038/s41598-020-72635-x.

19. Heppner, N., Reitmeier, S., Heddes, M., Merino, M.V., Schwartz, L., Dietrich, A., List, M., Gigl, M., Meng, C., van der Veen, D.R., et al. (2024). Diurnal rhythmicity of infant fecal microbiota and metabolites: A randomized controlled interventional trial with infant formula. Cell Host Microbe 32, 573–587.e5. 10.1016/j.chom.2024.02.015.

20. Laursen, M.F., Sakanaka, M., von Burg, N., Morbe, U., Andersen, D., Moll, J.M., Pekmez, C.T., Rivollier, A., Michaelsen, K.F., Mølgaard, C., et al. (2021). Bifidobacterium species associated with breastfeeding produce aromatic lactic acids in the infant gut. Nat. Microbiol. 2021 611 *6*, 1367–1382. 10.1038/s41564-021-00970-4.

21. Batta, V.K., Rao, S.C., and Patole, S.K. (2023). Bifidobacterium infantis as a probiotic in preterm infants: a systematic review and meta-analysis. Pediatr. Res. 94, 1887–1905. 10.1038/s41390-023-02716-w.

22. Ennis, D., Shmorak, S., Jantscher-Krenn, E., and Yassour, M. (2024). Longitudinal quantification of Bifidobacterium longum subsp. infantis reveals late colonization in the infant gut independent of maternal milk HMO composition. Nat. Commun. 2024 151 *15*, 894-. 10.1038/s41467-024-45209-y.

23. Shao, Y., Wang, S., Gichuki, B.M., Walson, J.L., Berkley, J.A., Lawley, T.D., Stares, M.D., Rozday, T.J., Kumar, N., Browne, H.P., et al. (2026). Genomic atlas of Bifidobacterium infantis and B. longum informs infant probiotic design. Cell 0. 10.1016/j.cell.2026.01.007.

24. Valles-Colomer, M., Blanco-Mfguez, A., Manghi, P., Asnicar, F., Dubois, L., Golzato, D., Armanini, F., Cumbo, F., Huang, K.D., Manara, S., et al. (2023). The person-to-person transmission landscape of the gut and oral microbiomes. Nature 614, 125–135. 10.1038/s41586-022-05620-1.

25. Catanzaro, J.R., Strauss, J.D., Bielecka, A., Porto, A.F., Lobo, F.M., Urban, A., Schofield, W.B., and Palm, N.W. (2019). IgA-deficient humans exhibit gut microbiota dysbiosis despite secretion of compensatory IgM. Sci. Rep. 9. 10.1038/s41598-019-49923-2.

26. Sender, R., Weiss, Y., Navon, Y., Milo, I., Azulay, N., Keren, L., Fuchs, S., Ben-Zvi, D., Noor, E., and Milo, R. (2023). The total mass, number, and distribution of immune cells in the human body. Proc. Natl. Acad. Sci. 120, e2308511120. 10.1073/pnas.2308511120.

27. Guo, J., Ren, C., Han, X., Huang, W., You, Y., and Zhan, J. (2021). Role of IgA in the early-life establishment of the gut microbiota and immunity: Implications for constructing a healthy start. Gut Microbes 13, 1908101. 10.1080/19490976.2021.1908101.

28. Huus, K.E., Petersen, C., and Finlay, B.B. (2021). Diversity and dynamism of IgA−microbiota interactions. Nat. Rev. Immunol. 2021 218 *21*, 514–525. 10.1038/s41577-021-00506-1.

29. Conrey, P.E., Denu, L., O’Boyle, K.C., Rozich, I., Green, J., Maslanka, J., Lubin, J.B., Duranova, T., Haltzman, B.L., Gianchetti, L., et al. (2023). IgA deficiency destabilizes homeostasis toward intestinal microbes and increases systemic immune dysregulation. Sci. Immunol. 8. 10.1126/SCIIMMUNOL.ADE2335.

30. Fadlallah, J., El Kafsi, H., Sterlin, D., Juste, C., Parizot, C., Dorgham, K., Autaa, G., Gouas, D., Almeida, M., Lepage, P., et al. (2018). Microbial ecology perturbation in human IgA deficiency. Sci. Transl. Med. 10. 10.1126/scitranslmed.aan1217.

31. Killeen, R.B., and Joseph, N.I. (2023). Selective IgA Deficiency. Allergy Clin. Immunol., 348–354. 10.1002/9781118609125.ch42.

32. Swain, S., Selmi, C., Gershwin, M.E., and Teuber, S.S. (2019). The clinical implications of selective IgA deficiency. J. Transl. Autoimmun. 2. 10.1016/j.jtauto.2019.100025.

33. Benjamini, Y., and Hochberg, Y. (1995). Controlling the False Discovery Rate: A Practical and Powerful Approach to Multiple Testing. J. R. Stat. Soc. Ser. B Stat. Methodol. 57, 289–300. 10.1111/J.2517-6161.1995.TB02031.X.

34. Kruskal, W.H., and Wallis, W.A. (1952). Use of Ranks in One-Criterion Variance Analysis. J. Am. Stat. Assoc. 47, 583–621. 10.1080/01621459.1952.10483441.

35. Gutierrez, M.J., Nino, G., Restrepo-Gualteros, S., Mondell, E., Chorvinsky, E., Bhattacharya, S., Bera, B.S., Welham, A., Hong, X., and Wang, X. (2024). Purine degradation pathway metabolites at birth and the risk of lower respiratory tract infections in infancy. ERJ open Res. 10. 10.1183/23120541.00693-2023.

36. Manghi, P., Antonello, G., Schiffer, L., Golzato, D., Wokaty, A., Beghini, F., Mirzayi, C., Long, K., Gravel-Pucillo, K., Piccinno, G., et al. (2025). Meta-analysis of 22,710 human microbiome metagenomes defines an oral-to-gut microbial enrichment score and associations with host health and disease. Nat. Commun. 2025 171 *17*, 196-. 10.1038/s41467-025-66888-1.

37. Carlino, N., Blanco-Míguez, A., Punčochář, M., Mengoni, C., Pinto, F., Tatti, A., Manghi, P., Armanini, F., Avagliano, M., Barcenilla, C., et al. (2024). Unexplored microbial diversity from 2,500 food metagenomes and links with the human microbiome. Cell 187, 5775–5795.e15. 10.1016/j.cell.2024.07.039.

38. Blanco-Mfguez, A., Beghini, F., Cumbo, F., McIver, L.J., Thompson, K.N., Zolfo, M., Manghi, P., Dubois, L., Huang, K.D., Thomas, A.M., et al. (2023). Extending and improving metagenomic taxonomic profiling with uncharacterized species using MetaPhlAn 4. Nat. Biotechnol. 41, 1633–1644. 10.1038/s41587-023-01688-w.

39. Xu, J., Duar, R.M., Quah, B., Gong, M., Tin, F., Chan, P., Sim, C.K., Tan, K.H., Chong, Y.S., Gluckman, P.D., et al. (2024). Delayed colonization of Bifidobacterium spp. and low prevalence of B. infantis among infants of Asian ancestry born in Singapore: insights from the GUSTO cohort study. Front. Pediatr. 12. 10.3389/FPED.2024.1421051.

40. Frese, S.A., Hutton, A.A., Contreras, L.N., Shaw, C.A., Palumbo, M.C., Casaburi, G., Xu, G., Davis, J.C.C., Lebrilla, C.B., Henrick, B.M., et al. (2017). Persistence of Supplemented Bifidobacterium longum subsp. infantis EVC001 in Breastfed Infants. mSphere 2. 10.1128/msphere.00501-17.

41. O’Brien, C.E., Meier, A.K., Cernioglo, K., Mitchell, R.D., Casaburi, G., Frese, S.A., Henrick, B.M., Underwood, M.A., and Smilowitz, J.T. (2022). Early probiotic supplementation with B. infantis in breastfed infants leads to persistent colonization at 1 year. Pediatr. Res. 91, 627–636. 10.1038/s41390-020-01350-0.

42. Bargheet, A., Hovde Bø, G., Andrea Klokkhammer Hetland, M., Langeland, N., Klingenberg, C., Kucha, V., Pettersen Correspondence, rová, Justine, M., John Moyo, S., Høyland Lohr, I., et al. (2026). Metabolic reprogramming of the infant gut by bifidobacteria-based probiotics drives exclusion of antibiotic-resistant pathobionts. Cell Reports Med. 7, 102752. 10.1016/J.XCRM.2026.102752.

43. Yu, D., Pei, Z., Chen, Y., Wang, H., Xiao, Y., Zhang, H., Chen, W., and Lu, W. (2023). Bifidobacterium longum subsp. infantis as widespread bacteriocin gene clusters carrier stands out among the Bifidobacterium. Appl. Environ. Microbiol. 89. 10.1128/aem.00979-23.

44. Kim, M., Qie, Y., Park, J., and Kim, C.H. (2016). Gut Microbial Metabolites Fuel Host Antibody Responses. Cell Host Microbe 20, 202–214. 10.1016/j.chom.2016.07.001.

45. Arzamasov, A.A., Rodionov, D.A., Hibberd, M.C., Guruge, J.L., Kent, J.E., Kazanov, M.D., Leyn, S.A., Elane, M.L., Sejane, K., Furst, A., et al. (2025). Integrative genomic reconstruction reveals heterogeneity in carbohydrate utilization across human gut bifidobacteria. Nat. Microbiol. 10, 2031–2047. 10.1038/s41564-025-02056-x.

46. Taft, D.H., Lewis, Z.T., Nguyen, N., Ho, S., Masarweh, C., Dunne-Castagna, V., Tancredi, D.J., Huda, M.N., Stephensen, C.B., Hinde, K., et al. (2022). Bifidobacterium Species Colonization in Infancy: A Global Cross-Sectional Comparison by Population History of Breastfeeding. Nutrients 14. 10.3390/NU14071423.

47. James, K., Motherway, M.O.C., Bottacini, F., and Van Sinderen, D. (2016). Bifidobacterium breve UCC2003 metabolises the human milk oligosaccharides lacto-N-tetraose and lacto-N-neo-tetraose through overlapping, yet distinct pathways. Sci. Rep. 6. 10.1038/srep38560.

48. LoCascio, R.G., Niiionuevo, M.R., Kronewitter, S.R., Freeman, S.L., German, J.B., Lebrilla, C.B., and Mills, D.A. (2009). A versatile and scalable strategy for glycoprofiling bifidobacterial consumption of human milk oligosaccharides. Microb. Biotechnol. 2, 333–342. 10.1111/j.1751-7915.2008.00072.x.

49. LoCascio, R.G., Desai, P., Sela, D.A., Weimer, B., and Mills, D.A. (2010). Broad conservation of milk utilization genes in Bifidobacterium longum subsp. infantis as revealed by comparative genomic hybridization. Appl. Environ. Microbiol. 76, 7373–7381. 10.1128/AEM.00675-10.

50. Underwood, M.A., German, J.B., Lebrilla, C.B., and Mills, D.A. (2015). Bifidobacterium longum subspecies infantis: champion colonizer of the infant gut. Pediatr. Res. 77, 229–235. 10.1038/pr.2014.156.

51. Zivkovic, A.M., German, J.B., Lebrilla, C.B., and Mills, D.A. (2011). Human milk glycobiome and its impact on the infant gastrointestinal microbiota. Proc. Natl. Acad. Sci. U. S. A. 108 *Suppl*, 4653–4658. 10.1073/pnas.1000083107.

52. Bu, K., Scherzi, T., Cantor, A., Teng, A.A., Pablo, J. V, Shandling, A.D., Randall, A.Z., Davis, E., Jackson, C., John Looney, R., et al. (2026). Gut microbial IgA coating in infants with traditional farming lifestyle and urban infants with allergic outcomes. Front. Immunol. 17, 1793302. 10.3389/FIMMU.2026.1793302.

53. Olm, M.R., Spencer, S.P., Takeuchi, T., Silva, E.L., and Sonnenburg, J.L. (2025). Metagenomic immunoglobulin sequencing reveals IgA coating of microbial strains in the healthy human gut. Nat. Microbiol. 10, 112–125. 10.1038/S41564-024-01887-4.

54. Reitmeier, S., Kiessling, S., Neuhaus, K., and Haller, D. (2020). Comparing Circadian Rhythmicity in the Human Gut Microbiome. STAR Protoc. 1. 10.1016/j.xpro.2020.100148.

55. Villette, R., Autaa, G., Hind, S., Holm, J.B., Moreno-Sabater, A., and Larsen, M. (2021). Refinement of 16S rRNA gene analysis for low biomass biospecimens. Sci. Reports 2021 111 11, 10741-. 10.1038/s41598-021-90226-2.

56. Babraham Bioinformatics - Trim Galore! https://www.bioinformatics.babraham.ac.uk/projects/trim_galore/.

57. Beghini, F., McIver, L.J., Blanco-Mfguez, A., Dubois, L., Asnicar, F., Maharjan, S., Mailyan, A., Manghi, P., Scholz, M., Thomas, A.M., et al. (2021). Integrating taxonomic, functional, and strain-level profiling of diverse microbial communities with bioBakery 3. Elife 10. 10.7554/eLife.65088.

58. Truong, D.T., Tett, A., Pasolli, E., Huttenhower, C., and Segata, N. (2017). Microbial strain-level population structure & genetic diversity from metagenomes. Genome Res. 27, 626–638. 10.1101/gr.216242.116.

59. Stamatakis, A. (2014). RAxML version 8: a tool for phylogenetic analysis and post-analysis of large phylogenies. Bioinformatics 30, 1312–1313. 10.1093/bioinformatics/btu033.

60. Letunic, I., and Bork, P. (2024). Interactive Tree of Life (iTOL) v6: recent updates to the phylogenetic tree display and annotation tool. Nucleic Acids Res. 52, W78–W82. 10.1093/nar/gkae268.

61. Asnicar, F., Weingart, G., Tickle, T.L., Huttenhower, C., and Segata, N. (2015). Compact graphical representation of phylogenetic data and metadata with GraPhlAn. PeerJ 2015, e1029. 10.7717/peerj.1029.

62. Caspi, R., Altman, T., Billington, R., Dreher, K., Foerster, H., Fulcher, C.A., Holland, T.A., Keseler, I.M., Kothari, A., Kubo, A., et al. (2014). The MetaCyc database of metabolic pathways and enzymes and the BioCyc collection of Pathway/Genome Databases. Nucleic Acids Res. 42, D459–D471. 10.1093/NAR/GKT1103.

63. Li, D., Liu, C.M., Luo, R., Sadakane, K., and Lam, T.W. (2015). MEGAHIT: an ultra-fast single-node solution for large and complex metagenomics assembly via succinct de Bruijn graph. Bioinformatics 31, 1674–1676. 10.1093/bioinformatics/btv033.

64. Hyatt, D., Chen, G.L., LoCascio, P.F., Land, M.L., Larimer, F.W., and Hauser, L.J. (2010). Prodigal: prokaryotic gene recognition and translation initiation site identification. BMC Bioinformatics 11. 10.1186/1471-2105-11-119.

65. Bazanella, M., Maier, T. V., Clavel, T., Lagkouvardos, I., Lucio, M., Maldonado-Gomez, M.X., Autran, C., Walter, J., Bode, L., Schmitt-Kopplin, P., et al. (2017). Randomized controlled trial on the impact of early-life intervention with bifidobacteria on the healthy infant fecal microbiota and metabolome. Am. J. Clin. Nutr. 106, 1274–1286. 10.3945/ajcn.117.157529.

66. Li, W., and Godzik, A. (2006). Cd-hit: a fast program for clustering and comparing large sets of protein or nucleotide sequences. Bioinformatics 22, 1658–1659. 10.1093/bioinformatics/btl158.

67. Oiiate, F.P., Le Chatelier, E., Almeida, M., Cervino, A.C.L., Gauthier, F., Magoules, F., Ehrlich, S.D., and Pichaud, M. (2019). MSPminer: abundance-based reconstitution of microbial pan-genomes from shotgun metagenomic data. Bioinformatics 35, 1544–1552. 10.1093/bioinformatics/bty830.

68. Chaumeil, P.A., Mussig, A.J., Hugenholtz, P., and Parks, D.H. (2019). GTDB-Tk: a toolkit to classify genomes with the Genome Taxonomy Database. Bioinformatics 36, 1925–1927. 10.1093/bioinformatics/btz848.

69. Jin, S., Cenier, A., Wetzel, D., Arefaine, B., Moreno-Gonzalez, M., Stamouli, M., Mohamad, M., Lupatsii, M., Rfos, E., Lee, S., et al. (2025). Microbial oral-gut translocation in advanced chronic liver disease is linked to exacerbation of intestinal barrier dysfunction and hepatic fibrosis. bioRxiv, 2025.09.23.677032. 10.1101/2025.09.23.677032.

70. Jones, P., Binns, D., Chang, H.Y., Fraser, M., Li, W., McAnulla, C., McWilliam, H., Maslen, J., Mitchell, A., Nuka, G., et al. (2014). InterProScan 5: genome-scale protein function classification. Bioinformatics 30, 1236–1240. 10.1093/bioinformatics/btu031.

71. Molder, F., Jablonski, K.P., Letcher, B., Hall, M.B., Tomkins-Tinch, C.H., Sochat, V., Forster, J., Lee, S., Twardziok, S.O., Kanitz, A., et al. (2021). Sustainable data analysis with Snakemake. F1000Research 10, 33. 10.12688/f1000research.29032.2.

72. Jin, S., Cenier, A., Eisenhard, L., Rfos, E., Wetzel, D., Cho, W.Y., Lesker, T.R., Eisen, T., Schorlemmer, S., Gaska, A., et al. (2025). Gene-centric metagenomic analyses reveal microbiome functional insights into diseases. bioRxiv, 2025.09.29.679262. 10.1101/2025.09.29.679262.

73. Oksanen, J., Simpson, G.L., Blanchet, F.G., Kindt, R., Legendre, P., Minchin, P.R., O’Hara, R.B., Solymos, P., Stevens, M.H.H., Szoecs, E., et al. (2025). vegan: Community Ecology Package, 10.32614/CRAN.package.vegan.

74. R Core Team (2025). R: A Language and Environment for Statistical Computing.

75. Martinez Arbizu, P. (2017). pairwiseAdonis: Pairwise Multilevel Comparison using Adonis.

76. Dinno, A. (2026). Dunn’s Test of Multiple Comparisons Using Rank Sums [R package dunn.test version 1.4.0]. CRAN Contrib. Packag. 10.32614/CRAN.PACKAGE.DUNN.TEST.

77. Wickham, H., Francois, R., Henry, L., Müller, K., and Vaughan, D. (2023). dplyr: A Grammar of Data Manipulation, 10.32614/CRAN.package.dplyr.

78. Wickham, H. (2016). ggplot2: Elegant Graphics for Data Analysis (Springer-Verlag New York).

79. Harris, C.R., Millman, K.J., van der Walt, S.J., Gommers, R., Virtanen, P., Cournapeau, D., Wieser, E., Taylor, J., Berg, S., Smith, N.J., et al. (2020). Array programming with NumPy. Nat. 2020 5857825 585, 357–362. 10.1038/s41586-020-2649-2.

80. Hunter, J.D. (2007). Matplotlib: A 2D graphics environment. Comput. Sci. Eng. 9, 90–95. 10.1109/MCSE.2007.55.

81. Waskom, M. (2021). seaborn: statistical data visualization. J. Open Source Softw. 6, 3021. 10.21105/joss.03021.

82. pandas development team, T. (2020). pandas-dev/pandas: Pandas at Zenodo, 10.5281/zenodo.3509134.

83. Virtanen, P., Gommers, R., Oliphant, T.E., Haberland, M., Reddy, T., Cournapeau, D., Burovski, E., Peterson, P., Weckesser, W., Bright, J., et al. (2020). SciPy 1.0: fundamental algorithms for scientific computing in Python. Nat. Methods 2020 173 *17*, 261–272. 10.1038/s41592-019-0686-2.

84. Charlier, F., Weber, M., Izak, D., Harkin, E., Magnus, M., Lalli, J., Fresnais, L., Chan, M., Markov, N., Amsalem, O., et al. (2022). Statannotations at Zenodo, 10.5281/zenodo.7213391.

85. Mann, H.B., and Whitney, D.R. (1947). On a Test of Whether one of Two Random Variables is Stochastically Larger than the Other. Ann. Math. Stat. 18, 50–60. 10.1214/aoms/1177730491.

86. Mallick, H., Rahnavard, A., McIver, L.J., Ma, S., Zhang, Y., Nguyen, L.H., Tickle, T.L., Weingart, G., Ren, B., Schwager, E.H., et al. (2021). Multivariable association discovery in population-scale meta-omics studies. PLOS Comput. Biol. 17, e1009442. 10.1371/journal.pcbi.1009442.

87. Slowikowski, K. (2024). ggrepel: Automatically Position Non-Overlapping Text Labels with “ggplot2,” 10.32614/CRAN.package.ggrepel.

88. Wilke, C.O., and Wiernik, B.M. (2022). ggtext: Improved Text Rendering Support for “ggplot2,” 10.32614/CRAN.package.ggtext.

